# Using FoxP2 to Distinguish Direct and Indirect Basal Ganglia Pathways for Vocal Learning in Songbirds

**DOI:** 10.1101/2024.12.08.627403

**Authors:** Aditi Jagannathan, Mira Nigudkar, Sarah W Bottjer

**Affiliations:** Section of Neurobiology, University of Southern California, Los Angeles, CA 90089 USA

**Keywords:** basal ganglia, thalamus, FoxP2, DARPP-32, songbird, vocal learning pallidal neurons

## Abstract

The cortico-basal ganglia pathways that mediate vocal learning in zebra finches (*Taeniopygia guttata*) are localized in parallel circuits formed by CORE and SHELL subregions. These circuits traverse a specialized region of the basal ganglia essential for vocal learning (Area X), which includes intermixed striatal and pallidal neurons. The pallidal neurons within Area X exhibit analogs of mammalian direct and indirect pathways that may have opposing effects and thereby increase or inhibit thalamic activity respectively. Direct pallidal neurons of Area X send projections to the medial portion of the dorsolateral anterior thalamic nucleus (DLM), whereas indirect pallidal neurons form intrinsic connections onto DLM-projecting neurons. Expression of the transcription factor FoxP2 in the basal ganglia is necessary for normal vocal learning and production in both humans and songbirds. We used tract-tracing techniques to label direct pallidal Area X→DLM projection neurons and immunohistochemical techniques to label neurons expressing the transcription factor FoxP2 in adult and juvenile male zebra finches. Our results showed that DLM-projecting neurons did not express FoxP2 in either adults or juveniles. Measurements of nuclear sizes revealed a population of large neurons that expressed FoxP2 but were not retrogradely-labeled from DLM. A putative marker of striatal neurons (DARPP-32) did not co-localize with FoxP2 in many of these large neurons, suggesting that they form a class of indirect pallidal neurons. These findings offer FoxP2 as a possible marker for indirect pallidal neurons and support the existence of different subpopulations of neurons that correspond to direct and indirect pathways within Area X.

## 1. INTRODUCTION

The basal ganglia in songbirds includes a specialized region essential for vocal learning and behavior called Area X (Fig. 1a), which receives input from two sensorimotor cortical regions that also regulate song-related functions: HVC and LMAN (Bottjer et al., 1989; Nixdorf-Bergweiler et al., 1995; Nottebohm et al., 1976; Sanchez-Valpuesta et al., 2019; Scharff & Nottebohm, 1991; Sohrabji et al., 1990; Trusel et al., 2025; Vates & Nottebohm, 1995). Area X sends inhibitory projections to the thalamic nucleus DLM, which projects back to LMAN (Luo et al., 2001; Luo & Perkel, 1999b; Nixdorf-Bergweiler et al., 1995; Vates & Nottebohm, 1995). The Area X→DLM→LMAN pathway consists of CORE and SHELL subregions which form parallel topographically organized recurrent circuits (Fig. 1a) (Bottjer et al., 1989; Gale et al., 2008; Iyengar et al., 1999; Johnson & Bottjer, 1992; Johnson et al., 1995; Luo et al., 2001; Person et al., 2008; Pinaud et al., 2007). Both CORE and SHELL loops are essential for vocal learning in juvenile birds (Achiro & Bottjer, 2013; Achiro et al., 2017; Bottjer & Altenau, 2010). In addition, Area X receives cerebellar inputs via thalamic nuclei adjacent to DLM (Nicholson et al., 2018; Person et al., 2008; Pidoux et al., 2018).

**Figure 1.**
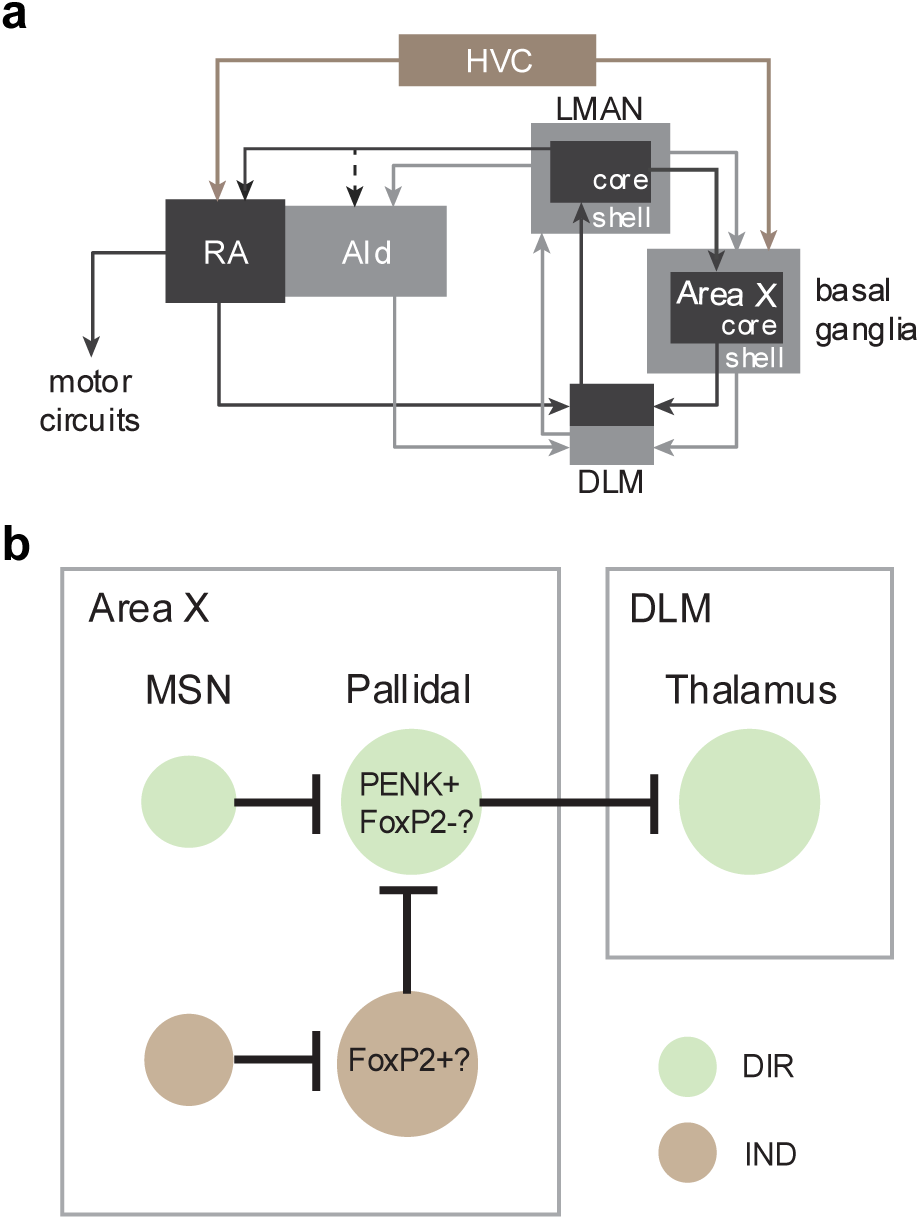
Schematics of cortico-basal ganglia circuits in zebra finches. (a) Simplified schematic of cortico-basal ganglia circuits that mediate vocal learning in songbirds. The recurrent pathway from LMAN→Area X→DLM comprises CORE and SHELL subregions that form independent parallel loops. (b) Schematic of putative direct (DIR) and indirect (IND) pathways from Area X to the thalamus (DLM). Abbreviations: RA, robust nucleus of the arcopallium; AId, dorsal intermediate arcopallium; LMAN, lateral magnocellular nucleus of the anterior nidopallium; DLM, medial portion of the dorsolateral anterior thalamic nucleus, HVC, high vocal center; MSN, medium spiny neuron.

The output of the CORE pathway from LMAN to vocal motor cortex (RA) drives song production in juvenile birds (Aronov et al., 2008; Bottjer et al., 1984; Scharff & Nottebohm, 1991). In contrast, the output of LMAN-SHELL projects to a region of motor cortex (AId) adjacent to RA that lacks direct motor targets; AId neurons make a strong projection to a dorsal thalamic zone which forms a transcortical loop that converges with information from Area X in DLM (Fig. 1a). In addition, AId neurons project to additional regions including substantia nigra compacta/ventral tegmental area (which projects to Area X), lateral hypothalamus, and the medial spiriform nucleus (which projects to the cerebellum) (Bottjer et al., 2000; Foster et al., 1997). Lesions of AId disrupt accurate imitation of the tutor song but do not impact vocal motor output in juvenile songbirds (Bottjer & Altenau, 2010). Therefore, SHELL circuitry does not drive vocal motor output, but may play a cognitive-limbic role in evaluating vocal output. The organization of CORE and SHELL loops is highly similar to sensorimotor and associative-limbic circuits of mammalian basal ganglia (Graybiel, 2008; Redgrave et al., 2010; Yin & Knowlton, 2006).

Area X contains analogs of both direct and indirect basal ganglia pathways (Farries et al., 2005; Goldberg et al., 2010). In the direct (DIR) pathway, pallidal neurons form calyceal inhibitory synapses on DLM neurons (Fig. 1b) (Carrillo & Doupe, 2004; Gale et al., 2008; Luo & Perkel, 1999a; Person et al., 2008; Reiner et al., 2004). A separate population of local pallidal neurons in Area X makes axonal contacts with DIR DLM-projecting neurons, forming an indirect (IND) pathway. These local neurons express GAD, and DLM-projecting neurons are surrounded by inhibitory GAD+ processes (Luo & Perkel, 1999b), suggesting that this non-projecting pallidal population forms intrinsic inhibitory projections onto DIR neurons of Area X (Fig. 1b).

Because dopamine receptors do not provide a clear marker for direct versus indirect neurons in songbirds (Kubikova et al., 2010), the transcription factor FoxP2 may serve as one possible marker to distinguish between direct and indirect pallidal neurons in Area X. FoxP2 is essential for human vocal development, with mutations leading to speech disorders such as childhood apraxia (den Hoed & Fisher, 2020). Likewise, disruptions of the FoxP2 gene in songbirds impair vocal learning and production and induce abnormal morphology in medium spiny neurons of Area X (Day et al., 2019; Haesler et al., 2007; Heston & White, 2015; Jarrell et al., 2025; Murugan et al., 2013; Shi et al., 2018; Xiao et al., 2021). Single nucleus RNA sequencing showed that a cluster of cells in Area X identified as pallidal formed two subgroups: one expressed the gene for enkephalin (ENK) and the other expressed the gene for FoxP2 (Xiao et al., 2021). Because ENK is expressed in DLM-projecting pallidal neurons in songbirds (Carrillo & Doupe, 2004), this pattern suggested that the direct neurons that project to DLM would not express FoxP2, while the indirect non-projecting pallidal neurons would be FoxP2+ (Fig. 1b). To test this idea, we used tract-tracing techniques to label direct pallidal Area X→DLM projection neurons and immunohistochemical techniques to label neurons expressing FoxP2. In addition, we measured the nuclear size of FoxP2+ nuclei to identify categories of different-sized cells and tested whether neurons with large FoxP2+ nuclei (potential pallidal neurons) lacked expression of DARPP-32, a putative marker of striatal neurons.

## 2. MATERIALS AND METHODS

### 2.1 Subjects

All animal procedures were performed in accordance with national regulatory guidelines (Public Health Service Policy), and followed protocols approved by the Animal Care and Use Committee at the University of Southern California. Zebra finches (*Taeniopygia guttata*) were bred in group aviaries in our colony and remained with their parents until at least 43-47 days post-hatch (dph), which is beyond the age of adequate tutor song exposure and social experience (Böhner, 1983, 1990; Catchpole & Slater, 1995; Clayton, 1987; Eales, 1985; Immelmann, 1969; Mann & Slater, 1995; Roper & Zann, 2006). Birds were moved to sex-specific holding cages in the colony after ∼45-50 dph. Intracranial injections of a fluorescent tracer were made in four adult male birds >90 dph (age range: 91-182, n = 4) and three juvenile male birds 45 ± 2 dph (n = 3). Following surgery, birds were placed in individual holding cages located in the main colony for three days before being perfused (see below).

### 2.2 Alexa Fluor 488 Injections

Birds were anesthetized with 1.5-1.7% isoflurane (inhalation) and placed in a stereotaxic apparatus. Stereotaxic coordinates were used to target the location of the thalamic nucleus DLM (dorsolateral nucleus of the medial thalamus; Fig. 1). The head angle was set as follows: we measured the depth of the skull at the bifurcation of the mid-sagittal sinus and at the midline 5 mm anterior to the bifurcation; the head angle was adjusted so that the difference in depth between these two locations was 2.2 ± 0.1. DLM injection coordinates were approximately 0.6 to 0.8 mm anterior and 0.9 to 1.1 mm lateral to the bifurcation of the mid-sagittal sinus at a depth of 4.5 mm in both juvenile and adult birds. Birds received unilateral left iontophoretic injections of dextran labeled with Alexa Fluor 488 (MW 10,000, 10% in sterile-filtered PBS, ThermoFisher #D-22910) into DLM by applying pulses of current through a silver wire (3-5 µA, 6 sec on/6 sec off) for 20 minutes through a glass pipette (20-30 μm outer diameter). Injections sometimes extended beyond the borders of DLM, but we analyzed retrograde label only within Area X-CORE and SHELL regions. Birds were anesthetized with an injection of 0.04 - 0.06 mL 5% Euthasol 72 hours following surgery and perfused transcardially with 0.7% saline followed by 4% periodate-lysine-paraformaldehyde (PLP) fixative (4% para in 0.1 M PB with 0.01 M sodium periodate and 0.1 M L-lysine; pH 7.8). Brains were removed and post-fixed in 4% paraformaldehyde overnight before being cryo-protected in 25% buffered sucrose solution with 0.005% sodium azide. Brains were frozen sectioned at a thickness of 40 µm in the coronal plane; sections were serially collected in PBS into separate series as described below.

### 2.3 Immunohistochemistry

#### 2.3.1 Double Label

Sections from four adult birds and one juvenile bird were labeled with retrograde Alexa Fluor 488 dye from DLM and stained using two different antibodies to FoxP2 (Table 2). Three separate series were processed as follows: Series 1 was stained using rabbit polyclonal anti-FoxP2 (Abcam Cat# ab16046, RRID:AB_2107107), Series 2 was stained using goat polyclonal anti-FoxP2 (Abcam Cat# ab1307, RRID:AB_1268914), and Series 3 was counterstained with thionin for Nissl and coverslipped with Permount (Fisher Scientific). The ab16046 antibody was generated using a synthetic peptide corresponding to human FoxP2 aa 700 to the C-terminus conjugated to keyhole limpet haemocyanin (specificity has been described previously, Campbell et al., 2009). The ab1307 antibody was raised against a synthetic peptide corresponding to amino acids 703-715 (C-terminal) of human FoxP2 (specificity has been described previously, Thompson et al., 2013).

Sections were stained using standard immunohistochemistry techniques. Following three washes in 0.02 M PBS, Series 1 and 2 were blocked in 5% Normal Goat Serum or 5% Normal Rabbit Serum, respectively, for 30-40 minutes and then incubated in the primary antibody (1:1,000) overnight at room temperature. Sections were again triple-washed with PBS and incubated in the appropriate secondary antibody (anti-rabbit-Alexa 594, 1:200, ThermoFisher, #A-11012, made in goat for Series 1; anti-goat-Alexa 594, 1:200, ThermoFisher, #A-11080, made in rabbit for Series 2) for 1 hour. Sections of both series were given three final washes in PBS with the last wash containing 1 μL DAPI (1:10,000, gift from Dr. Jason Junge) before being mounted onto gelatin-coated glass slides. Sections were allowed to air dry before being coverslipped with Prolong Diamond Antifade (ThermoFisher, #P36970).

After processing these 5 birds (four adults and one juvenile), we analyzed the size of neuronal nuclei and observed a continuous range of FoxP2+ nuclear sizes (shown in Fig. 5 below). We had expected to observe a dichotomous distribution that would segregate medium spiny and pallidal neurons (see Results). Since the continuous distribution of nuclear size clearly made this strategy untenable, we decided to test DARPP-32 as an additional marker, which selectively labels medium spiny neurons in mammals, for the remaining two juvenile birds as explained in the next section.

#### 2.3.2 Triple Label

DARPP-32 is a phosphoprotein that is expressed in virtually all medium spiny neurons of mammals (Ouimet et al., 1998), although some studies have reported it to be immunohistochemically detectable in only 30-60% of spiny neurons in birds (Reiner et al., 2004; Reiner et al., 1998; Singh & Iyengar, 2019). We attempted to test the identity of cells labeled by FoxP2 by performing triple labeling for FoxP2, DARPP-32, and the retrograde label from DLM in two juvenile birds. During prior analysis of double-labeled tissue, we observed that the anti-FoxP2 used in Series 1 (ab16046) resulted in a higher density of labeled cells compared to the anti-FoxP2 used in Series 2 (ab1307). We therefore created a cocktail of rabbit polyclonal anti-FoxP2 (ab16046, 1:1,000) and mouse monoclonal anti-DARPP-32 [1:30,000, C24-6a, gift from Dr. Hugh C. Hemmings, specificity validated in Hemmings and Greengard (1986) and see Durstewitz et al. (1998) for pigeon data]; alternate sections were stained for Nissl. The same blocking and primary staining procedures outlined for Series 1 above were followed before incubating sections in a secondary antibody cocktail containing anti-rabbit Alexa 594 (1:200, ThermoFisher, #A11012, made in goat) and anti-mouse Alexa 647 (1:200, ThermoFisher, #A-21235, made in goat). All markers used in this study are outlined in Table 1.

**Table 1.** Fluorescent label, type of cellular stain, confocal microscopy excitation/emission wavelengths, and color used in figures for each label in the experiment.

| Label | Type of stain | Excitation/emission<br>(nm) | Color in figures |
| --- | --- | --- | --- |
| **DAPI** | Nuclear | 410/497 | Blue |
| **DLM-projecting**<br>Alexa Fluor 488 | Cytoplasmic | 500/580 | Green |
| **FoxP2**<br>Alexa Fluor 594 | Nuclear | 589/651 | Red |
| **DARPP-32**<br>Alexa Fluor 647 | Cytoplasmic | 653/755 | Magenta |

### 2.4 Data Analysis

#### 2.4.1 Overlap Analysis of FoxP2 and Retrogradely-Labeled Neurons in Area X

Patterns of fluorescent label were analyzed in all adult (n = 4) and juvenile (n = 3) birds. Images of Medial Striatum, which includes Area X, were taken with a 40x objective using ZEN imaging software on a confocal microscope (Zeiss LSM 780 inverted microscope or Zeiss LSM 880 inverted microscope) at the USC Translational Imaging Center. The excitation and emission wavelength settings used to image each marker are given in Table 1. The tilescan feature of ZEN was used to acquire multiple individual image fields (“tiles”) at adjacent XY positions. To collect depth information, the ZEN Z-stack feature was used to image multiple Z positions (“Z-steps”) for each tile. Each tile within a tissue section included 8-12 Z-steps that were taken approximately 3-4 µm apart. All tiles from a single section were stitched together using the Imaris Stitcher application (Oxford Instruments); the Z-steps were reconstructed into a 3D projection of the section in Imaris image analysis software. 3D projections of individual or multiple fluorescent channels were used in all figures except Figure 7, which was taken at a single Z-step (see below, 2.4.4). All 3D projections were visualized using the maximum intensity projection display mode in Imaris, which presents the voxel with maximum intensity from all Z-steps in the viewing plane.

3D projections were analyzed for overlap between neurons retrogradely-labeled from DLM and neurons expressing FoxP2. Both antibodies against FoxP2 (ab16046 and ab1307) were used in all four adults and one juvenile for overlap analysis. Because qualitative examination in these five birds revealed that anti-FoxP2 16046 produced a higher incidence of labeled cells than anti-FoxP2 1307, we used ab16046 to stain the remaining two juvenile birds. As a result, in juvenile overlap analysis, only one bird was labeled for both FoxP2 antibodies and the remaining two birds were labeled with the 16046 antibody only (Table 2). Retrogradely-labeled cells (adults: n = 469; juveniles: n = 87) were classified as being within either Area X-CORE or Area X-SHELL. The borders of X-CORE were outlined by DAPI staining. Because the borders of X-SHELL are not visible based on cytoarchitecture, we used previous data from our lab in which X-SHELL was defined by afferent inputs from LMAN-SHELL to define X-SHELL conservatively as 500 µm extending from the medial, ventromedial, and ventrolateral edges of Area X-CORE (Iyengar et al., 1999). Neurons were assigned to a “Border” category if the boundary between Area X-CORE and SHELL was not easily identifiable by DAPI staining. Each retrogradely-labeled cell was visually inspected at every Z-step for signs of double-label with FoxP2.

**Table 2.** Age and number of DLM-projecting DIR neurons in Area X-CORE and Area X-SHELL. . †These columns list the number of DIR DLM-projecting neurons for each bird. ‡This column indicates the number of DIR neurons included in the overlap analysis based on staining from each antibody against FoxP2. If DAPI staining was poor on a given tissue section and could not be used to define the precise borders of Area X-CORE, the DLM-projecting direct neurons were placed in the “Border” column. See text for further details.

| Bird ID | Age (days post-hatch) | # of DIR neurons in Area X-CORE† | # of DIR neurons in Area X-SHELL† | # of DIR neurons on border of Area X-CORE and Area X-SHELL† | # of DIR neurons from sections stained with ab16046/ab1307‡ |
| --- | --- | --- | --- | --- | --- |
| Db51 | 167 | 98 | 30 | 10 | 67/71 |
| R27 | 182 | 58 | 55 | 20 | 60/73 |
| Db69 | 104 | 39 | 31 | 26 | 43/53 |
| R54 | 91 | 47 | 52 | 3 | 46/56 |
|  |  | **242** | **168** | **59** | **216/253**<br>**Adult n = 469** |
| Gy124 | 46 | 27 | 15 | 18 | 60/na |
| Gy48 | 44 | 9 | 8 | 4 | 21/na |
| O63 | 45 | 2 | 3 | 1 | 2/4 |
|  |  | **38** | **26** | **23** | **83/4**<br>**Juvenile n = 87** |

#### 2.4.2 Size Analysis of Retrogradely-Labeled Somata in Area X

Size analysis of Area X neurons retrogradely-labeled from DLM was performed on all four adult birds (n = 100 neurons chosen randomly for analysis out of a total of 469 retrogradely-labeled neurons) and all three juvenile birds (n = 87 neurons). Using the Surfaces auto-detection feature in Imaris, 3D models of labeled cells were created based on the Z-intervals described above in 2.4.1 (3-4 µm). Cells were detected using automatic absolute intensity thresholding on the Alexa Fluor 488 channel and smoothed with a surface detail of 0.50 µm to ensure close fitting to the staining. Once the Surfaces feature created 3D models of labeled cells, we manually inspected each neuron and adjusted each surface to include only the cell body, excluding the primary dendrites.

We measured the diameter of these 3D models of retrogradely-labeled neurons by using the average of two measures from Imaris Surfaces: the longest diameter of each soma (referred to as Object-Oriented Bounding Box Length C) and the perpendicular diameter (Object-Oriented Bounding Box Length B); we refer to this measure as the average diameter. This measure is consistent with previous studies in songbirds that reported the average of the major and minor axis of somata as the diameter (Luo & Perkel, 1999b; Reiner et al., 2004). The average diameter values of each Alexa 488-labeled surface were exported to Microsoft Excel for further analysis.

#### 2.4.3 Size Analysis of FoxP2-Positive Nuclei in Area X

Size analysis of FoxP2+ nuclei in Area X-CORE and SHELL was performed on three adult and two juvenile birds. One adult and one juvenile were excluded from analysis because high background staining made it difficult for the Surfaces auto-detection feature of Imaris to accurately distinguish individual nuclei. Only sections stained for anti-FoxP2 16046 were examined in the FoxP2+ size analysis. Because the full depth profile of smaller FoxP2+ neurons was not represented in the Z-stack intervals used in the overlap analysis of DIR pallidal cells described above, two to five tissue sections per bird were reimaged with smaller Z-stack intervals (0.46 - 0.47 µm apart). This interval ensured that smaller cell types, such as medium spiny neurons, were included in the analysis. The boundaries of Area X-CORE in each section were determined using DAPI staining and divided into 212.55 µm x 212.55 µm fields. Images of two randomly selected fields per section were taken at a magnification of 40x with the updated Z-stack intervals. Two 212.55 µm x 212.55 µm fields per section were also taken for Area X-SHELL: one image was taken ventromedial and the other ventrolateral to the boundaries of Area X-CORE.

We determined the size of FoxP2+ nuclei using the Surfaces auto-detection feature of Imaris on the Alexa Fluor 594 channel; surfaces were smoothed with a surface detail of 0.50 µm (Fig. 2). Adjoining surfaces were split based on morphology with a seed point diameter of 3.20 µm to account for the smallest possible striatal medium spiny neuron diameter (Carrillo & Doupe, 2004). Cells were then filtered based on area (>25 µm^2^) and distance from the XYZ border (>0.5 µm) to remove cells that were only partially included in the section. Each 3D projection generated by the Surfaces feature was visually inspected to see if the 3D model matched the FoxP2 staining; if the model’s outline was not accurate, the surface was deleted. Additionally, all neurons that exhibited a fusiform appearance were excluded from analysis; these cells are likely migratory young neurons (Alvarez-Buylla & Nottebohm, 1988). Because larger FoxP2+ nuclei appeared to be more intensely stained (see below), we took protective measures during image acquisition and analysis to avoid saturation. During image acquisition, we adjusted laser power and detector gain to optimal values using the Range Indicator display tool. During analysis of each section, we also carefully reviewed the histogram of FoxP2+ nuclei Surfaces to ensure that intensities were not clustered to the maximum intensity value (which would suggest oversaturation).

**Figure 2.**
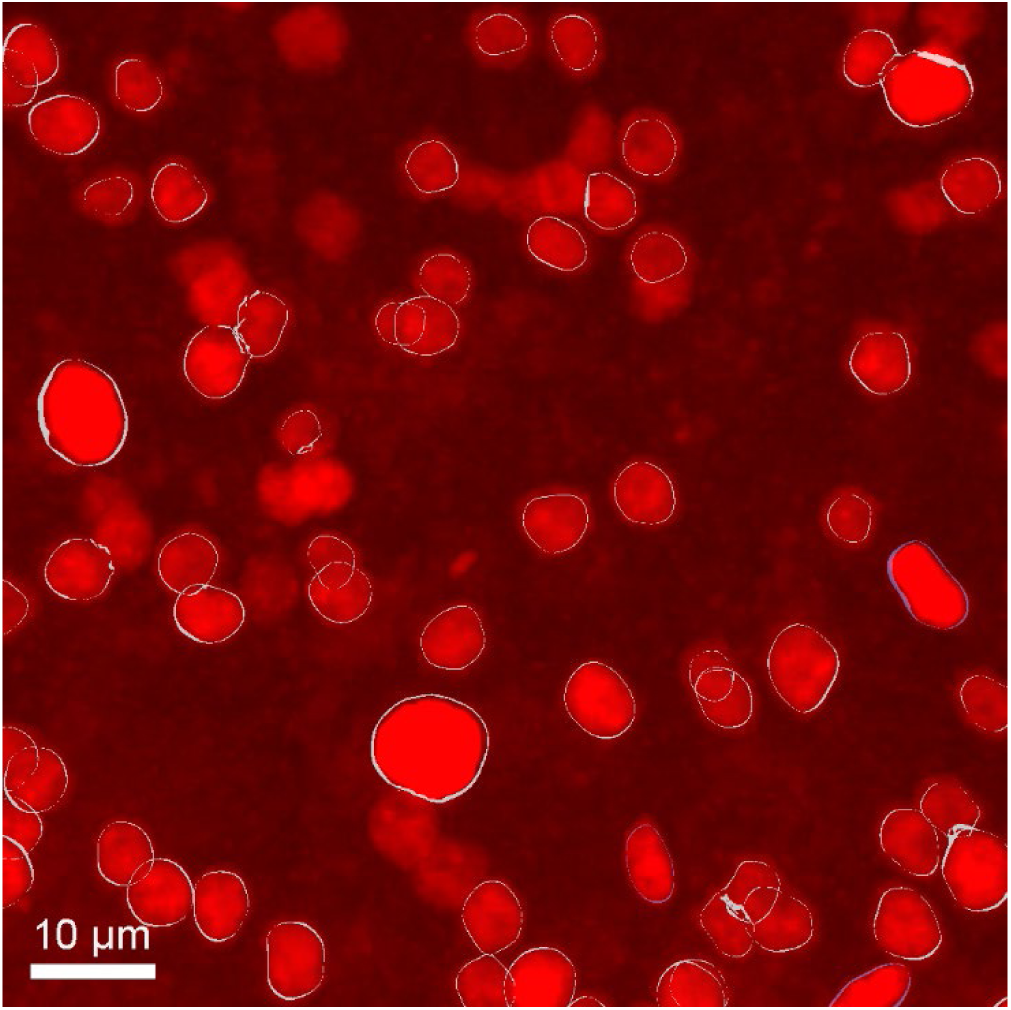
Surfaces auto-detection feature was used to measure size of FoxP2+ nuclei. The size of FoxP2+ nuclei was measured using the Surfaces auto-detection feature of Imaris; confocal image shows white outlines around FoxP2+ nuclei. See text for further details.

In adults, 3,953 and 1,535 FoxP2+ nuclei were obtained from Area X-CORE and Area X-SHELL, respectively. In Area X-CORE and Area X-SHELL of juveniles, 1,813 and 835 FoxP2+ nuclei were analyzed. Measurements of average diameter were calculated as explained in section 2.4.2 and exported to Microsoft Excel for further analysis.

#### 2.4.4 Overlap Analysis of Large FoxP2+ Neurons with DARPP-32 in juvenile birds

As detailed below in Results, we observed a population of large FoxP2+ neurons that were not retrogradely-labeled from DLM, representing possible IND pallidal neurons in Area X. However, because FoxP2 was expressed in nuclei of different sizes along a continuous distribution (see below, Fig. 5), it was not possible to use this measure to segregate non-projecting FoxP2+ pallidal neurons from smaller FoxP2+ cells based on size. We decided to use nuclear diameter as a means of segregating putative IND neurons given that their nuclear diameter should be similar to that of DIR DLM-projecting neurons. Based on this idea, we first measured the nuclear diameter of 50 retrogradely-labeled DIR neurons (which were FoxP2-negative) from one adult bird in which DAPI staining was particularly good. The full depth profile of each DIR neuron was inspected using the original Z-steps of 3-4 µm (see above, 2.4.1) and the Z-step that included the largest visible nucleus was selected. The Measurement Points feature of Imaris was then used to determine the longest diameter and the perpendicular diameter of the nucleus at the selected Z-step. The mean of these two values was calculated in Microsoft Excel and reported as the average nuclear diameter for DIR neurons. It should be noted that the average diameter values may represent an underestimate since there was no way to confirm that each cell was imaged at the Z-plane where the nucleus was the largest. The mean diameter of these adult DIR pallidal nuclei was 7.95 ± 1.01 µm and did not differ between neurons in Area X-CORE and SHELL (p = 0.2941; unpaired t-test). We therefore established an initial cutoff of 8 µm to separate putative IND pallidal neurons from smaller medium spiny neurons based on nuclear size. While this measurement was established from adult data (due to optimal DAPI staining), we applied the nuclear size cutoff to juveniles because the Alexa 488+ DIR somal size did not differ between adult and juvenile songbirds (see below).

Neurons with large (≥8 um) FoxP2+ nuclei in two juvenile birds with staining for the putative striatal marker DARPP-32 were analyzed for overlap to assess whether these cells were pallidal in nature. We examined all FoxP2+ neurons with an average nuclear diameter ≥8 µm (n = 96; putative IND pallidal neurons) across Area X-CORE and SHELL. Each neuron was visually inspected at every Z position (at a step size of 0.46-0.47 µm) for signs of FoxP2/DARPP-32 co-expression. For purposes of illustration, the voxel-based Colocalization feature in Imaris was used to display voxels that co-expressed both FoxP2 and DARPP-32 channels. The threshold for FoxP2 and DARPP-32 channel colocalization was set at the single Z-step where the FoxP2+ nucleus appeared the largest and was then applied to the entire Z-stack to visualize double label (see Fig. 6).

#### 2.4.5 Statistical Tests

All continuous data were tested for normality using D’Agostino-Pearson tests in GraphPad Prism software. An unpaired t-test was used to compare mean diameters of adult DIR pallidal nuclear size between Area X regions. A two-factor ANOVA (age x region) was used to test differences in the size of retrogradely labeled somata in juveniles versus adults and X-CORE versus X-SHELL using GraphPad Prism, as these data were normally distributed.

Planned comparisons were tested using Fisher’s Least Significant Difference (LSD) test and Benjamini–Hochberg corrections were applied for multiple comparisons (Benjamini & Hochberg, 1995). Differences in the size of FoxP2+ nuclei were non-normal, and were tested using a two-factor Gamma GLM (Generalized Linear Model) with a log link in R. The glm function built into R was used to fit the GLM, and the emmeans package was used to calculate pairwise comparisons. R version 4.4.1 was used for all analyses. Differences in the proportions of neurons were assessed for statistical significance using the Chi-square test. When there were only two possible outcomes, a binomial test was used to analyze differences in the expected and observed proportions. The expected proportion of DIR neurons that co-expressed FoxP2 was set to 0.5 to represent random chance. Measures of central tendency for normally and non-normally distributed data are reported as mean ± standard deviation and median with first and third quartiles (Table 3), respectively.

**Table 3.** Median diameter, Q1, and Q3 values reported for size in µm (top), nucleus count (middle), and relative percentage (bottom) for large (≥8 µm) and small (<8 µm) FoxP2+ nuclei in Area X-CORE and Area X-SHELL of adults and juveniles.

| | Large FoxP2+ ($\geq 8 \mu\text{m}$) | | Small FoxP2+ ($< 8 \mu\text{m}$) | |
| --- | --- | --- | --- | --- |
|  | Area X-CORE | Area X-SHELL | Area X-CORE | Area X-SHELL |
| **Adult** | 8.35 (8.15, 8.77)<br>n = 102<br>2.58% | 8.17 (8.09, 8.47)<br>n = 40<br>2.61% | 5.31 (4.66, 6.02)<br>n = 3851<br>97.42% | 5.41 (4.74, 6.15)<br>n = 1495<br>97.39% |
| **Juvenile** | 8.43 (8.23, 8.70)<br>n = 48<br>2.65% | 8.37 (8.15, 8.87)<br>n = 48<br>5.75% | 5.85 (5.15, 6.61)<br>n = 1765<br>97.35% | 5.70 (4.82, 6.60)<br>n = 787<br>94.25% |

#### 2.4.6 Ethical statement

The authors confirm that no part of this manuscript has been published elsewhere. No GenAI tools were used to develop any portion of the text of this manuscript.

#### 2.4.7 Data Availability

The data that support the findings of this study are available from the corresponding author upon reasonable request.

## 3. RESULTS

### 3.1 Direct DLM-projecting neurons in Area X do not express FoxP2 in adult or juvenile songbirds

We injected dextran-Alexa 488 into DLM in order to retrogradely label thalamus-projecting (DIR) neurons within CORE and SHELL regions of Area X (see Methods; Fig. 3a, 4a). DAPI staining was used to identify the outline of Area X-CORE (Fig. 3b, 4b); Area X-SHELL was defined conservatively as extending 500 µm from the medial, ventromedial, and ventrolateral edges of Area X-CORE (Iyengar et al., 1999; see Methods). Our injections revealed a topographically organized pattern of retrograde label in Area X-CORE and Area X-SHELL that was consistent with previous work (Iyengar et al., 1999; Johnson et al., 1995; Luo & Perkel, 1999b; Person et al., 2008) (data not shown). Some injections extended beyond the borders of DLM, resulting in labeled neurons outside of Area X-CORE and SHELL; we did not include these neurons in our analysis. The precise borders of Area X-CORE were poorly defined in some sections, and some retrogradely-labeled neurons were located very close to the CORE-SHELL border. In these cases, we assigned labeled neurons to the border region between Area X-CORE and Area X-SHELL.

**Figure 3.**
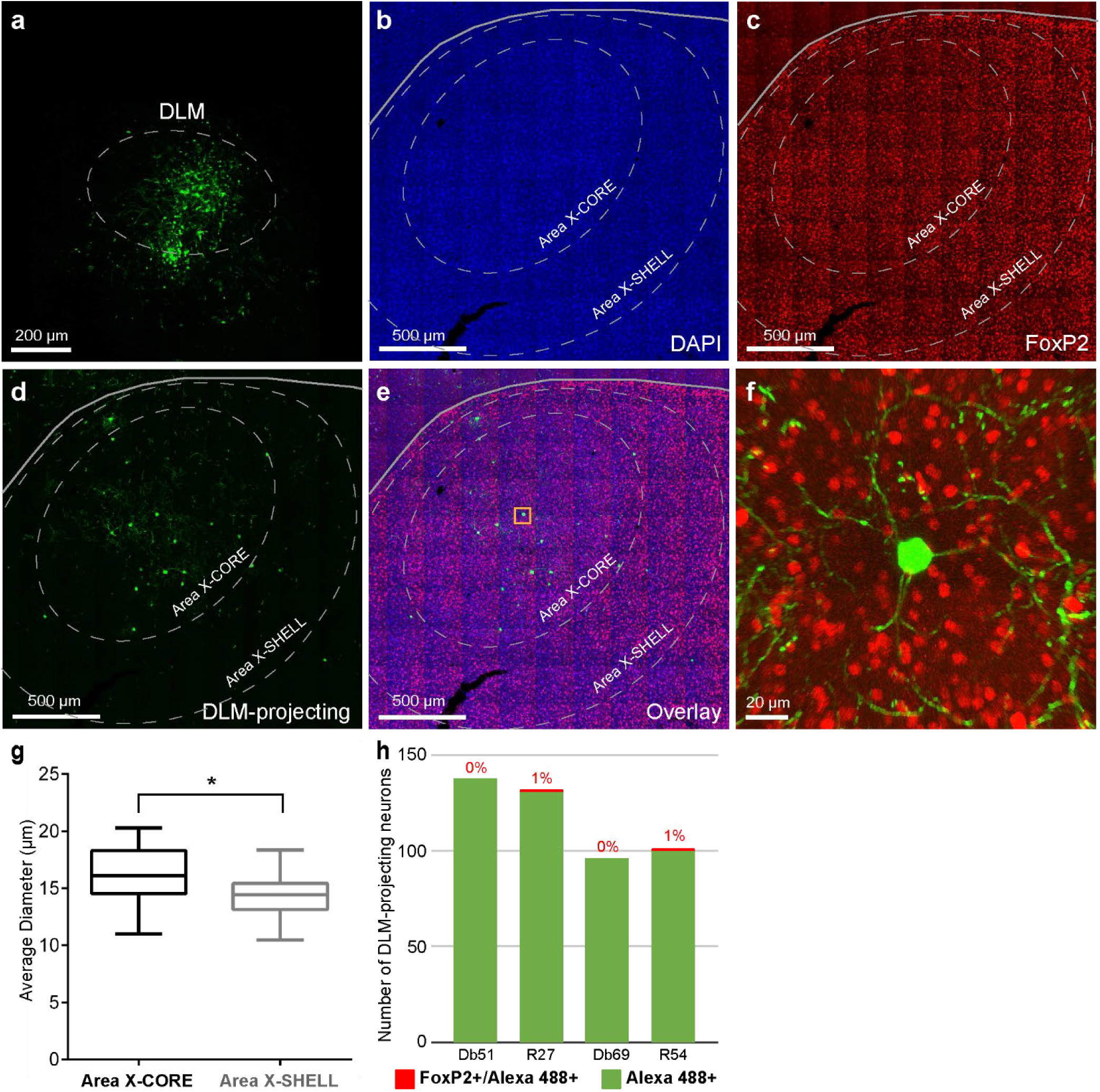
DIR neurons in Area X of adult zebra finches do not express FoxP2. (a) Example of fluorescent tracer injection into DLM (outlined by dashed line). Injection produced retrograde label in Area X-CORE and Area X-SHELL of an adult bird (d-f). Injection site extended beyond the ventral border of DLM, but we did not analyze labeled neurons outside of CORE and SHELL regions of Area X (see Methods). (b) DAPI staining outlines Area X-CORE (inner dashed circle) within Medial Striatum of basal ganglia (gray line marks the axon tract PSL, pallial-subpallial lamina, which separates striatum from cortex). Area X-SHELL (outer dashed circle) was conservatively defined as extending 500 µm from the medial, ventromedial, and ventrolateral edges of Area X-CORE. (c) Red FoxP2 staining (Alexa 594) using 16046 antibody and (d) green retrogradely-labeled cells from DLM (Alexa 488) in the same section as (b). (e) Overlay of DAPI, FoxP2, and DLM-projecting dyes. (f) Enlarged image of boxed region in (e) demonstrating that red FoxP2 label and green DLM-projecting retrograde label did not colocalize in a DIR neuron in Area X-CORE. (g) Box plots showing that the mean somal diameter of DIR DLM-projecting neurons in Area X-CORE (n = 54) was larger than those in Area X-SHELL (n = 35) (*p < 0.0004; Fisher’s LSD test with Benjamini-Hochberg correction). (h) Bar graph showing that 467/469 DIR neurons in Area X-CORE and SHELL of 4 adult birds did not express FoxP2 (p < 0.0001, binomial test). Percentages represent number of DIR neurons that expressed FoxP2 over total number of DIR neurons in each bird.

**Figure 4.**
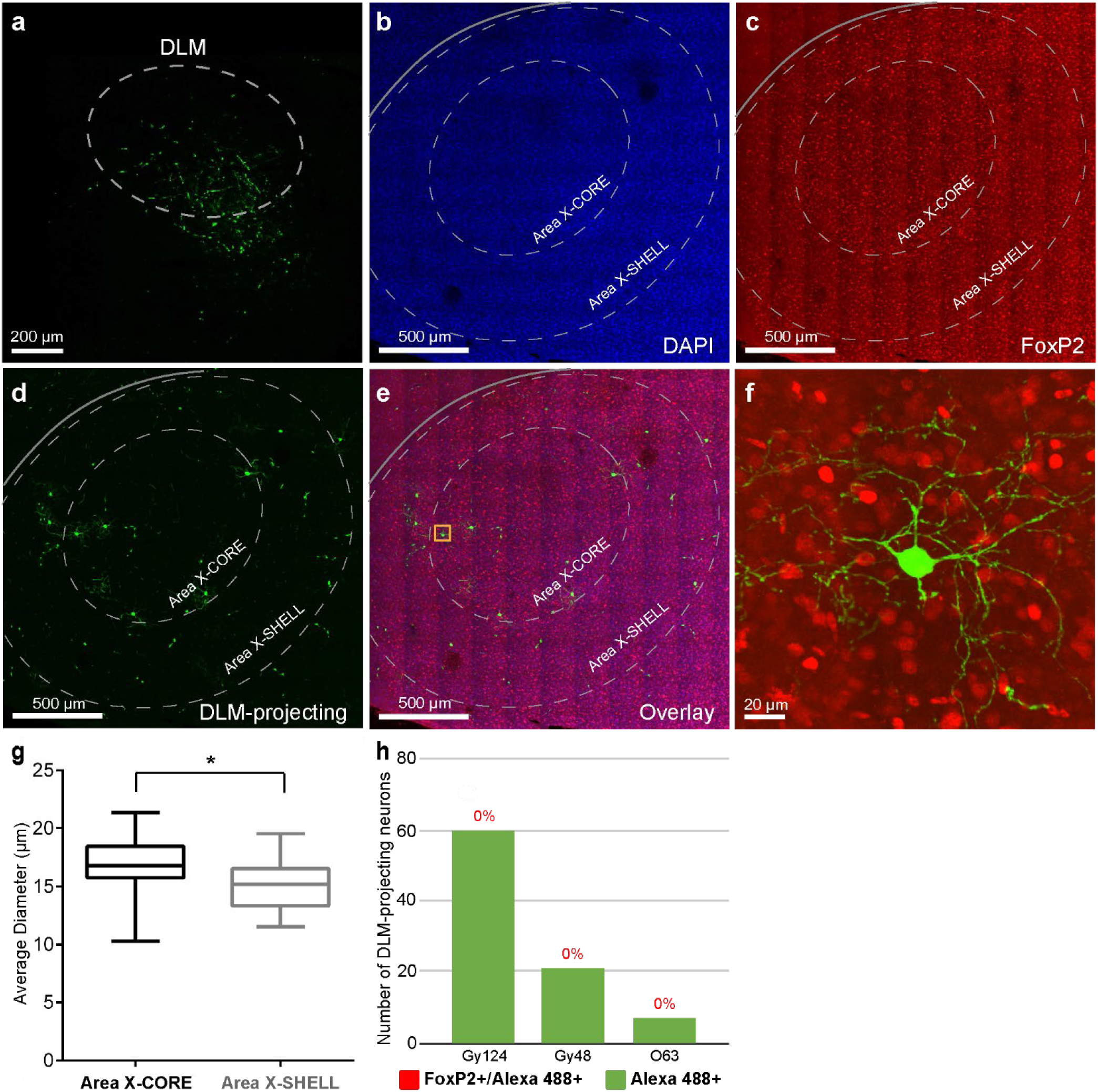
DIR neurons in Area X of juvenile songbirds do not express FoxP2. (a) Injection of a fluorescent tracer into DLM (outlined by dashed line) produced retrograde label in Area X-CORE and SHELL of a juvenile bird (d-f). (b) DAPI staining outlines Area X-CORE (inner dashed circle) within Medial Striatum (the border between striatum and cortex is demarcated by gray line, which corresponds to the axon tract PSL, pallial-subpallial lamina). As in adult birds, Area X-SHELL (outer dashed circle) was conservatively defined as extending 500 µm from the medial, ventromedial, and ventrolateral edges of Area X-CORE. (c) Red FoxP2 staining (Alexa 594) using 16046 antibody and (d) green DLM-projecting neurons (Alexa 488) in the same section as (b). (e) Overlay of DAPI, FoxP2, and DLM-projecting dyes. (f) Enlarged image of boxed region in (e) demonstrating that red FoxP2 label and green retrograde label in an Area X-CORE DIR neuron did not colocalize. (g) Box plot showing that the mean somal diameter of DIR DLM-projecting neurons in Area X-CORE (n = 38) was larger than those in Area X-SHELL (n = 26) (*p = 0.001; Fisher’s LSD test with Benjamini-Hochberg correction). (h) Bar graph detailing that no DIR neurons in three juvenile birds (n = 87) showed FoxP2 expression (p < 0.0001, binomial test). Percentages represent number of DIR neurons that expressed FoxP2 over total number of DIR neurons in each bird.

Sections were stained in two series by different antibodies for FoxP2 (anti-FoxP2 16046 or anti-FoxP2 1307, see Methods; Fig. 3c). Table 2 shows the number of DIR DLM-projecting neurons in Area X-CORE, Area X-SHELL, and the border between them in each bird, as well as the number of DIR neurons analyzed from each FoxP2 antibody. The somata and primary dendrites of Area X neurons that projected to DLM were robustly labeled by the retrogradely transported dextran-Alexa 488 (Fig. 3f). These primary dendrites were aspiny in nature, characteristic of pallidal neurons as described by Reiner et al. (2004). We analyzed a total of 469 retrogradely-labeled neurons in four adult male birds, including 242 in Area X-CORE, 168 in Area X-SHELL, and 59 in the border region (Fig. 3d-f). All retrogradely-labeled neurons were inspected for overlap with FoxP2 (see Methods). Of these 469 DLM-projecting neurons, only two neurons in Area X-CORE showed possible co-expression of Alexa 488 (DLM-projecting) and Alexa 594 (FoxP2) labels (p < 0.0001, binomial test) (Fig. 3h). We next examined whether the absence of FoxP2 expression in adult DLM-projecting DIR neurons was also true in juvenile zebra finches (n = 3; Fig. 4c, d). Eighty-seven retrogradely-labeled cells were analyzed for FoxP2 co-expression (Fig. 4e, f), including 38 in Area X-CORE, 26 in Area X-SHELL, and 23 in the border region between CORE and SHELL (Table 2). None of these juvenile DIR neurons showed expression of FoxP2 (Fig. 4h). Together, these data show that DIR Area X→DLM projection neurons do not express FoxP2 in either adult or 45-dph juvenile zebra finches.

To determine whether the somal size of DIR neurons differed between Area X-CORE versus Area X-SHELL, 100 DIR neurons from the 469 retrogradely-labeled neurons across four adult birds were randomly selected for size measurements. Of these, 54 were in Area X-CORE, 35 were in Area X-SHELL, and 11 were in the border region; the border region neurons were excluded from size analysis because they could not be definitively categorized into either subregion. The DIR somata in adult Area X-CORE had a mean diameter of 16.29 ± 2.17 µm (n = 54). In comparison, DIR somata in adult Area X-SHELL had a mean diameter of 14.39 ± 1.59 µm (n = 35) (Fig. 3g). The size of DLM-projecting somata in juvenile Area X-CORE (n = 38) was 17.05 ± 2.28 µm whereas the size of those in Area X-SHELL (n = 26) was15.13 ± 2.28 µm (Fig. 4g). A two-factor ANOVA (age x region) showed two main effects: somata in X-CORE were larger than those in X-SHELL (p < 0.0001) and juvenile somata were larger than those in adults (p = 0.034); the interaction was not significant (p = 0.977). Planned comparisons (Benjamini-Hochberg corrected) revealed that somata in X-CORE were larger than those in X-SHELL in both adults and juveniles (p < 0.0004 and p = 0.001, respectively). However, the difference in somal size between adults and juveniles was not significant in either X-CORE (p = 0.119) or X-SHELL (p = 0.176). This pattern shows that the somal size of DLM-projecting neurons is similar in adult and 45-dph juvenile birds, and is larger in Area X-CORE compared to X-SHELL in both juveniles and adults.

### 3.2 FoxP2+ nuclei vary in size and intensity

FoxP2+ nuclei in Area X exhibited a range of sizes in both adult (n = 3) and juvenile birds (n = 2) (Fig. 5a, c). Frequency histograms of size across all FoxP2+ nuclei detected using the 16046 antibody revealed a continuous, positively skewed distribution for adults and juveniles within both CORE and SHELL regions of Area X (Fig. 5b). In Area X-CORE, the average diameter of FoxP2-labeled nuclei ranged from 3.02 µm to 9.97 µm in adults and 3.21 to 10.14 µm in juveniles. In Area X-SHELL, the average diameter of labeled nuclei ranged from 3.21 µm to 9.61 µm in adults and 3.46 µm to 10.02 µm in juveniles.

**Figure 5.**
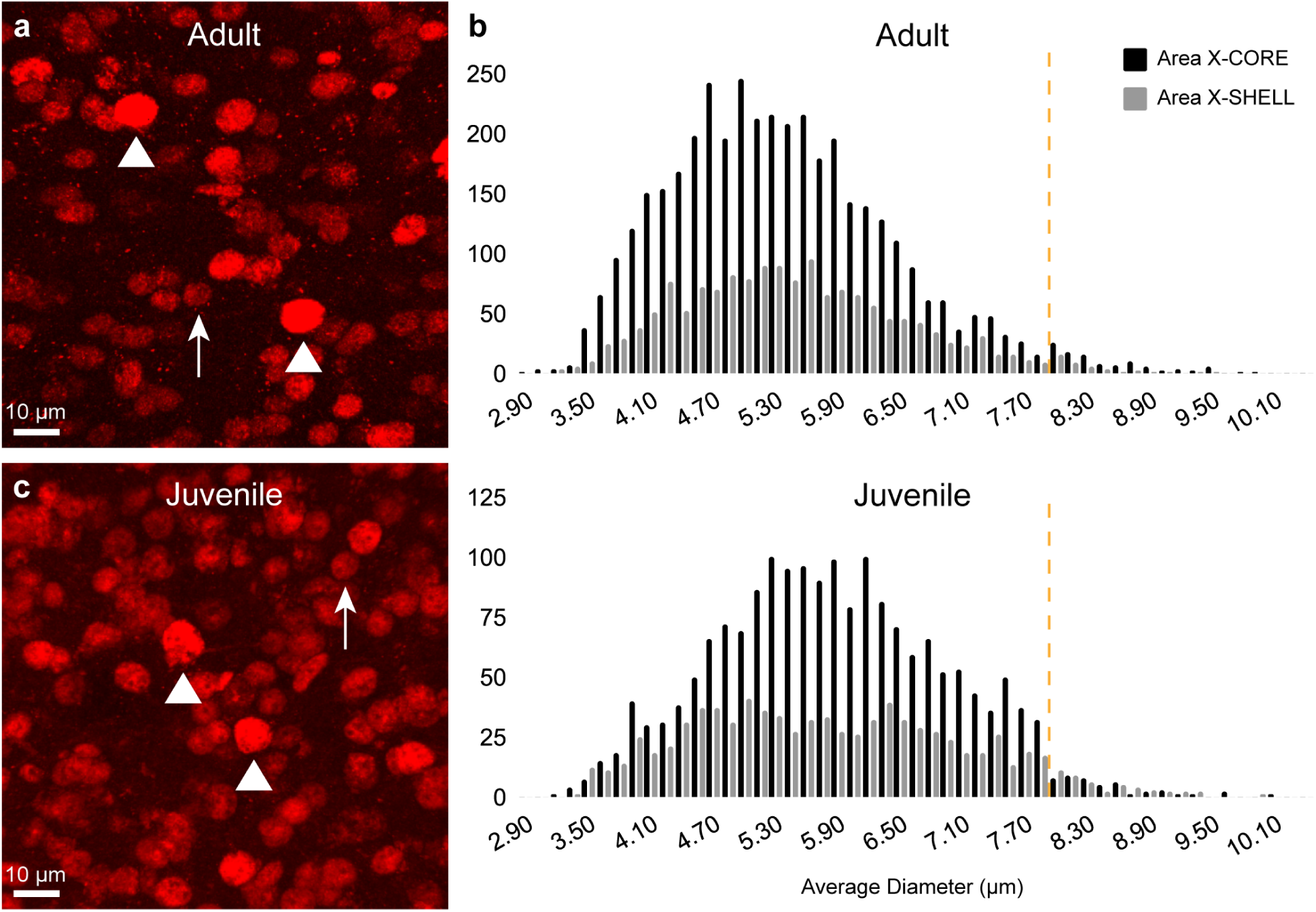
Average diameters of FoxP2+ nuclei in Area X-CORE and SHELL revealed continuous, positively skewed size distributions. FoxP2 staining distinguished sparsely distributed large nuclei (white arrowheads) and numerous small nuclei (white arrows) in Area X-CORE of (a) adult and (c) juvenile songbirds. This pattern was also present in Area X-SHELL (not shown). Most small FoxP2+ neurons were less intensely stained, whereas larger nuclei were more intensely stained. (b) Histograms showing average diameter of FoxP2+ nuclei in Area X-CORE (black, n = 3953) and SHELL (gray, n = 1535) of three adult birds (top) and two juvenile birds (bottom; CORE, black, n = 1813; SHELL, gray, n = 835). Only sections stained for anti-FoxP2 16046 were examined in the FoxP2+ size analysis (see Methods). Yellow dashed line indicates our initial cutoff value for large FoxP2+ neurons. Note the difference in scale of the y-axis between adults and juveniles.

Pallidal cells in the Medial Striatum of zebra finches, including Area X, are larger than striatal medium spiny neurons (Luo & Perkel, 1999b; Reiner et al., 2004), as is true in mammals. Reiner et al. (2004) previously reported that DLM-projecting pallidal neurons in zebra finches had average somal diameters of 12.4 ± 1.8 µm. In contrast, they found that the somata of medium spiny neurons, marked by the striatal cytoplasmic marker DARPP-32, had a diameter of 7.1 ± 1.1 µm. Interestingly, we did not observe a bimodal distribution in size suggested by this difference in diameter between medium spiny neurons and pallidal neurons (Fig. 5b), possibly due to the fact that FoxP2 is expressed not only in spiny neurons and pallidal-like neurons, but also oligodendrocytes and some types of interneurons in Area X (Haesler et al., 2004; Xiao et al., 2021). Because of the known size disparity between spiny and pallidal neurons across taxa, we hypothesized that most of the smaller FoxP2+ neurons corresponded to medium spiny neurons (Fig. 5a, c; white arrows), while the sparsely distributed larger FoxP2+ neurons corresponded to IND pallidal neurons (Fig. 5a, c; white arrowheads).

To gauge whether large FoxP2+ cells belonged to a distinct subpopulation, we first estimated the nuclear size of DLM-projecting pallidal neurons (see Methods). This analysis indicated that nuclei of DIR DLM-projecting neurons had a mean diameter of 7.95 ± 1.01 µm. We therefore established 8 µm as an initial cutoff for the average nuclear diameter of possible FoxP2+ indirect pallidal neurons. In Area X-CORE of adults, 97.42% of FoxP2+ nuclei had an average diameter <8 µm (referred to as “small FoxP2+ cells”); the median diameter of these nuclei was 5.31 µm (Table 3). In Area X-SHELL, small FoxP2+ nuclei accounted for 97.39% of the total cell count in adults with a median diameter of 5.41 µm. In juveniles, small FoxP2+ nuclei comprised 97.35% of the total cell count in Area X-CORE with a median diameter of 5.85 µm, whereas in Area X-SHELL, 94.25% were small FoxP2+ cells with a median diameter of 5.70 µm (Table 3). We tested differences in the size of small FoxP2+ nuclei using a two-factor (age x region) Gamma GLM with a log link (see Methods), which revealed main effects of age (juvenile nuclei larger than adult, p < 1 x 10^-16^) and region (p = 0.002), as well as their interaction (8.3 x 10^-6^). These results were confirmed by planned comparisons (Benjamini-Hochberg corrected), which showed that juvenile nuclei were larger than adults in both Area X-CORE (p < 0.0002) and Area X-SHELL (p < 0.0002). The interaction was due to the fact that small nuclei in juvenile X-CORE were larger than those in X-SHELL (p = 0.002), while small nuclei in adult X-SHELL were larger than in X-CORE (p = 0.002). Thus, small nuclei in juveniles were larger overall than those in adults, and juvenile nuclei were larger in X-CORE compared to X-SHELL whereas adult nuclei were larger in X-SHELL than in X-CORE.

We focused primarily on large FoxP2+ nuclei because they represent a possible population of IND pallidal neurons. In adult zebra finches, 2.58% of FoxP2+ nuclei in Area X-CORE fell above the cutoff of ≥8 µm with a median diameter of 8.35 µm (Table 3). In adult Area X-SHELL, 2.61% of FoxP2+ nuclei were large and had a median diameter of 8.17 µm. The proportion of large FoxP2+ nuclei did not differ between CORE and SHELL subregions (p = 0.957; Chi-square test). In juvenile songbirds, 2.65% and 5.75% of FoxP2+ nuclei were categorized as large (≥8 µm) in Area X-CORE and Area X-SHELL, respectively. This higher proportion of large FoxP2+ nuclei in juvenile Area X-SHELL compared to juvenile Area X-CORE was significant (p < 0.0001; Chi-square test). The proportion of large FoxP2+ nuclei did not differ across adults and juveniles in Area X-CORE (p = 0.8816; Chi-square test); however, Area X-SHELL of juveniles contained a greater proportion of large FoxP2+ nuclei compared to adult Area X-SHELL (p = 0.0001; Chi-square test), suggesting a developmental difference in the cellular makeup of this region. A two-factor (age x region) Gamma GLM with a log link showed that large nuclei in X-CORE were larger than those in X-SHELL (p = 0.038) and that the size of juvenile and adult nuclei did not differ (p = 0.572); the interaction was not significant (p = 0.265). Planned comparisons for region (Benjamini-Hochberg corrected) showed that large nuclei did not differ in size between X-CORE and X-SHELL in either adults (p = 0.075) or juveniles (p = 0.693). This pattern of results suggests a trend towards larger FoxP2+ nuclei in X-CORE compared to X-SHELL in adult birds only.

Interestingly, our qualitative observations suggested that these large nuclei were generally more intensely stained for FoxP2 than their smaller counterparts in both adults and juveniles (Fig. 5a, c). This observation aligns with data from Thompson et al. (2013) and Kosubek-Langer and Scharff (2020), which showed a bimodal distribution of relatively larger intensely-stained FoxP2+ nuclei and relatively smaller weakly-stained FoxP2+ nuclei in Area X. In summary, our size analysis revealed that a small percentage of FoxP2+ neurons in Area X had nuclei as large as those of pallidal DLM-projecting neurons and tended to show relatively intense FoxP2 labeling. This result is consistent with the sparse distribution of neurons in Area X of zebra finches identified by the pallidal marker LANT6 (Reiner et al., 2004).

### 3.3 Many large FoxP2+ neurons do not express DARPP-32 in juvenile birds

In order to further investigate whether the larger FoxP2+ neurons were pallidal in nature, we examined the expression of DARPP-32, which marks ∼50% or more of striatal spiny neurons in avian brain (Reiner et al., 2004; Reiner et al., 1998; Singh & Iyengar, 2019). DLM-projecting cells never expressed DARPP-32 in Area X-CORE or Area X-SHELL (Fig. 6a), consistent with a pallidal identity of DIR neurons. Therefore, if larger FoxP2+ neurons belong to an indirect pallidal class, we expected to observe a lack of overlap with DARPP-32 expression in this class of neurons also.

**Figure 6.**
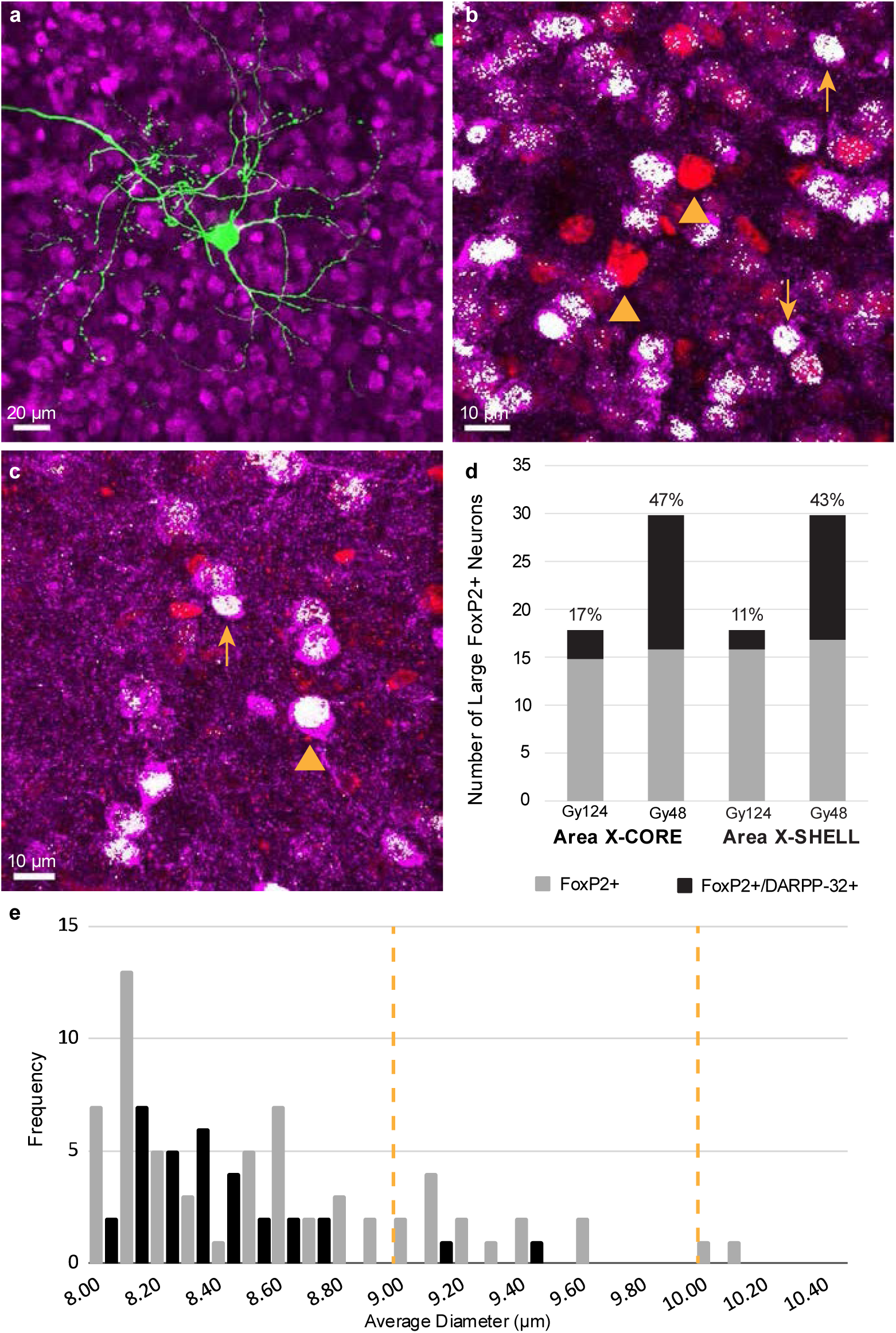
DARPP-32 was not expressed in DIR neurons and in many large putative IND neurons of juvenile songbirds. (a) Alexa 488-labeled DIR DLM-projecting neurons in juveniles (one shown from Area X-CORE) did not colocalize with magenta cytoplasmic DARPP-32 label. (b) Most large (≥8 μm) putative IND FoxP2+ nuclei (red) across both Area X-CORE and SHELL did not express DARPP-32 (magenta); two examples from Area-X CORE are indicated by yellow arrowheads. A majority of small (<8 μm) FoxP2+ nuclei expressed DARPP-32 (two examples indicated by yellow arrows). Colocalization of both proteins is depicted in white. (c) Some large (≥8 μm) FoxP2+ nuclei expressed DARPP-32 in both CORE and SHELL regions of Area X; yellow arrowhead indicates a large FoxP2+/DARPP-32+ neuron from Area-X SHELL. A nearby small (<8 μm; yellow arrow) FoxP2+ nucleus also expressed DARPP-32. White voxels indicating colocalization show that the FoxP2+ nuclei of the putative IND and medium spiny neuron take up a large percentage of the DARPP-32+ cytoplasm. (d) Bar graph detailing the number of large FoxP2+ neurons that show DARPP-32 expression in Area X-CORE (17/48 neurons) and Area X-SHELL (15/48 neurons) from two juvenile songbirds when a nuclear diameter cutoff of ≥8 µm was applied. Percentages represent number of large FoxP2+/DARPP32+ nuclei over total number of large FoxP2+ nuclei across CORE and SHELL subregions for each bird. (e) Histogram showing the average diameters of all large (≥8 µm) FoxP2+/DARPP-32- (gray; n = 64) and FoxP2+/DARPP-32+ (black; n = 32) neurons across Area X-CORE and SHELL. Yellow dashed lines indicate the cutoff values for large FoxP2+ neurons when increased by 1 and 2 standard deviations to 9.01 µm and 10.02 µm, respectively.

We analyzed all large FoxP2+ neurons with an average nuclear diameter ≥8 µm across both Area X-CORE and Area X-SHELL in two juvenile birds (n = 48 within each subregion, Table 3; see Methods). FoxP2 staining (red) was inspected for co-expression with DARPP-32 staining (magenta); white voxels depict colocalization of the two channels (Fig. 6b, c). Many large FoxP2+ neurons did not express DARPP-32 in either Area-X CORE (65%, 31/48) or Area X-SHELL (69%, 33/48); two examples from Area X-CORE are marked by yellow arrowheads in Figure 6b. However, a substantial number of larger neurons did co-express FoxP2 and DARPP-32 in both CORE and SHELL regions of Area X. That is, 35% (17/48) of large FoxP2+ neurons were double-labeled in Area X-CORE and 31% (15/48) were double-labeled in Area X-SHELL in these two juvenile birds (Fig. 6d). Figure 6c shows a large neuron in Area X-SHELL that expressed both FoxP2 and DARPP-32 (yellow arrowhead). This surprising result is consistent with the snRNA-seq findings of Xiao et al. (2021) in Area X that DARPP-32 is expressed in multiple cell types, including pallidal neurons.

Because our cutoff for large neurons represented a fairly rudimentary criterion, we decided to re-assess the incidence of FoxP2 and DARPP-32 co-expression in these 96 neurons when the cutoff value of 8.0 µm was increased by one or two standard deviations. Interestingly, when the cutoff value was increased by one standard deviation to 9.01 (SD = 1.01 µm), only 2% (1/48) of large FoxP2+ nuclei in Area X-CORE expressed DARPP-32. When the cutoff was raised by two standard deviations to 10.02, 0% (0/48) of large FoxP2+ nuclei in Area X-CORE were also DARPP-32+. Similarly, when the cutoff value was raised by one or two standard deviations only 2% (1/48) and 0% (0/48) of large FoxP2+ nuclei in Area X-SHELL expressed DARPP-32, respectively. Figure 6e shows the incidence of double labeling as a function of nuclear size collapsed across X-CORE and X-SHELL neurons (n = 96). This distribution shows that larger FoxP2-positive nuclei tend not to express DARPP-32.

We examined smaller neurons qualitatively, which revealed that a large majority expressed both FoxP2 and DARPP-32 as seen by the white voxels depicting colocalization (Fig. 6b, c; yellow arrows). Most of these likely represent medium spiny neurons, although it is possible that other small cell types such as some interneurons and glial cells also express FoxP2 in Area X (Xiao et al., 2021). The continuous distribution of sizes in FoxP2+ nuclei that we observed (Fig. 5b) is also consistent with the idea that DARPP-32 is not a selective marker for medium spiny neurons in zebra finch brain, a finding that is similar to results in pigeons (Reiner et al., 2004; Reiner et al., 1998; Singh & Iyengar, 2019).

Our qualitative observations of smaller neurons in Area X-CORE and SHELL also revealed that FoxP2+ nuclei of double-labeled neurons took up a large percentage of the cytoplasm labeled by DARPP-32, examples of which can be seen in Figures 6b and 6c (yellow arrows). These observations conform to the report of Ouimet et al. (1998) that DARPP-32+ medium spiny neurons of rat striatum have scant cytoplasm. Qualitative inspection also suggested that the nuclei of large FoxP2+/DARPP-32+ cells (including possible IND pallidal neurons) in Area X-CORE and SHELL follow a similar pattern by taking up a large percentage of the cytoplasm. In Figure 6c, the nuclear FoxP2 staining in a large FoxP2+ neuron (yellow arrowhead) takes up a large percentage of the cytoplasmic DARPP-32 staining, as seen by the white pixels indicating colocalization. Conversely, when we examined DIR retrogradely-labeled neurons, we observed that the DAPI+ nuclei encompassed a considerably smaller percentage of the soma. This observation was consistent across both Area X-CORE and SHELL regions. Figure 7 shows a DIR neuron in Area X-CORE in which the DAPI+ nucleus (blue) is surrounded by relatively more cytoplasm (green) compared to the large neuron in Figure 6c (yellow arrowhead). An adjacent FoxP2+ nucleus may represent an interneuron or a medium spiny neuron. These observations suggest that DIR and IND pallidal cells have a similar nuclear size but vary in the percentage of the soma that their nuclei encompass. DIR neurons may have a larger amount of cytoplasm in order to support a longer axon and/or other features (Al Ghamdi et al., 2009; Nolan et al., 2024; Tomasi et al., 2012).

**Figure 7.**
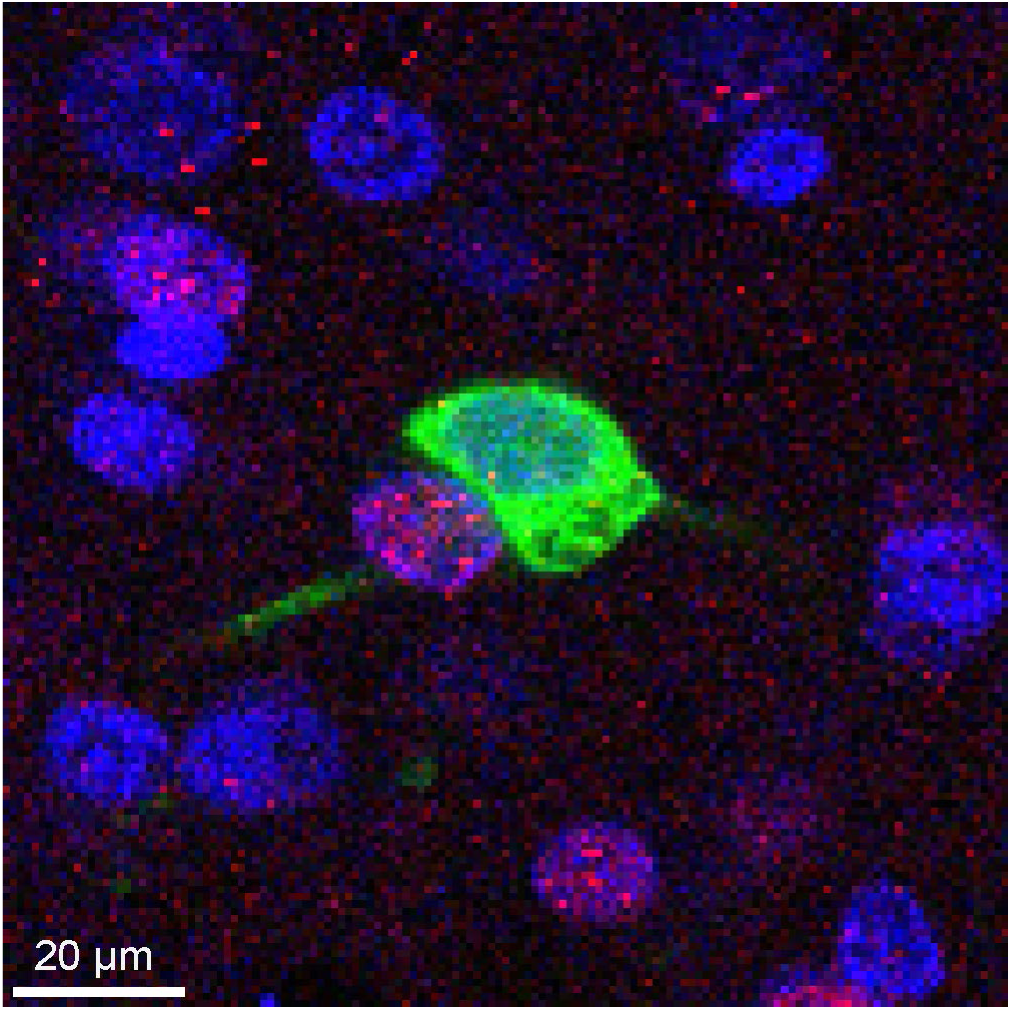
DLM-projecting DIR pallidal cell and adjacent FoxP2+ neuron have similar DAPI+ nuclear sizes. The nucleus of a green Alexa 488+ retrogradely-labeled cell is defined by blue DAPI nuclear stain in Area X-CORE of an adult bird. DAPI+ nucleus of DIR neuron encompasses a smaller percentage of the soma in comparison to putative IND large FoxP2+/DARPP-32+ cells (see Fig. 6c); this pattern was also present in Area X-SHELL (not shown). The adjacent red FoxP2+ nucleus may represent an interneuron or a medium spiny neuron.

## 4. DISCUSSION

Vocal learning is mediated in part by two parallel topographically organized cortico-basal ganglia circuits, CORE and SHELL, that form reentrant loops (Fig. 1a). Area X sends inhibitory projections to the thalamic nucleus DLM: Area X-CORE projects to the dorsolateral subregion of DLM (DLMdl) and Area X-SHELL projects to the ventromedial subregion of DLM (DLMvm) (Bottjer et al., 1989; Carrillo & Doupe, 2004; Iyengar et al., 1999; Johnson et al., 1995; Person et al., 2008). DLMdl and DLMvm send excitatory projections to LMAN-CORE and LMAN-SHELL respectively, forming recurrent pathways (Iyengar et al., 1999; Johnson et al., 1995; Luo et al., 2001; Pinaud et al., 2007). Thus, DLM receives information from independent Area X-CORE and SHELL pathways as well as convergent input from AId (Bottjer et al., 2000; Foster et al., 1997). Given the complexity of these connections, it remains unclear how motor production and vocal evaluation information are integrated in this circuitry (Iyengar et al., 1999; Johnson & Bottjer, 1992; Johnson et al., 1995; Paterson & Bottjer, 2017). Our goal in this study was to investigate specific patterns of connectivity underlying how vocal information is conveyed from Area X to DLM. A prior study using snRNA-seq showed that pallidal neurons in Area X express either FoxP2 or PENK (the precursor to enkephalin), but not both (Xiao et al., 2021). Given that enkephalin is the neurotransmitter of DIR DLM-projecting neurons (Carrillo & Doupe, 2004), this distinction suggested the hypothesis that DLM-projecting pallidal cells are FoxP2–/PENK+, while non-projecting pallidal cells are FoxP2+/PENK–. In this study, we tested FoxP2 expression as a possible marker for direct versus indirect pallidal neurons in Area X as a first step to distinguish how information may be differently conveyed via these two pathways from Area X→DLM.

Consistent with this hypothesis, we found that DIR DLM-projecting neurons in CORE and SHELL regions of Area X of both adult and juvenile songbirds lacked expression of FoxP2. Neurons with nuclei that did express FoxP2 exhibited various sizes and intensities in both CORE and SHELL regions of Area X. Qualitative inspection, along with nuclear size analysis, revealed a subpopulation of larger FoxP2+ neurons that appeared more intensely stained, consistent with findings from Thompson et al. (2013) and Kosubek-Langer and Scharff (2020). Coupled with the results from Xiao et al. (2021) regarding a lack of colocalization between FoxP2 and PENK in Area X, this pattern of results suggests that a group of larger, more intense FoxP2+ neurons represent a class of indirect pallidal neurons (Farries et al., 2005), while smaller FoxP2+ neurons represent spiny neurons and other cell types.

As a preliminary test of whether larger FoxP2+ (putative IND) neurons had a pallidal identity, we examined expression of the putative striatal neuron marker DARPP-32. Surprisingly, we discovered that 32/96 (33%) large (≥8 µm diameter) FoxP2+ neurons also expressed DARPP-32 in juvenile birds. Increasing our average size cutoff value by one or two standard deviations greatly reduced or eliminated the incidence of FoxP2 and DARPP-32 co-expression, suggesting perhaps that most IND pallidal neurons do not express DARPP-32. An alternative interpretation is that Area X contains diverse classes of FoxP2+ pallidal cells which have a larger range of nuclear sizes compared to DIR pallidal cells, including some that may be homologous to diverse classes of GPe pallidal neurons in mammals (Hegeman et al., 2016; Mastro et al., 2017; Spix et al., 2021; Vounatsos & Gittis, 2025). A lack of DARPP-32 expression has been reported in pallidal neurons of rodents and pigeons, although the exact antibody used has not always been specified, which may contribute to variable results (Anderson & Reiner, 1991; Reiner et al., 1998). FoxP2 does not colocalize with antibody markers for choline acetyltransferase (ChAT), nitric oxide (nNOS/SS/NPY), or parvalbumin (PV) interneurons in Area X (Haesler et al., 2004). DARPP-32 has also not been detected in cholinergic, PV+, or NPY+ striatal interneurons of rat or pigeon (Anderson & Reiner, 1991; Reiner et al., 1998). However, recent studies of the molecular profile of songbird and chicken brains have shown that a distinct class of GABAergic neurons derived from the LGE (lateral ganglionic eminence) expresses FoxP2 and occurs throughout the telencephalon (Colquitt et al., 2021; Zaremba et al., 2025). In mammals, this population is expressed primarily in striatum (spiny neuron) and ventral pallium (olfactory bulb and pallial amygdala), but in songbird Area X this class includes both spiny and pallidal neurons. In addition, the snRNA-seq analysis of Xiao et al. (2021) in adult Area X indicated that FoxP2 is expressed not only in pallidal neurons but also in interneurons (type unknown), oligodendrocytes, and endothelial cells, and that DARPP-32 (PPP1R1B) is expressed at a low level in some pallidal neurons. This latter finding is consistent with our observation that DARPP-32 and FoxP2 co-localized to a subset of pallidal neurons in juvenile birds.

Indeed, the growing complexity of cell types and connections in basal ganglia generally (Arber & Costa, 2022; Bostan et al., 2018; Courtney et al., 2023; Hegeman et al., 2016; Oorschot, 2016; Soares-Cunha et al., 2016; Spix et al., 2021; Vounatsos & Gittis, 2025), as well as the combined presence of spiny neurons, pallidal cells, interneurons, and possibly other neuronal cell types within Medial Striatum of avian brain, offer multiple possibilities for the identity of large FoxP2+/DARPP-32+ neurons in Area X. Mammalian GPe contains a heterogeneous assortment of neurons with diverse connections (Hegeman et al., 2016; Spix et al., 2021; Vounatsos & Gittis, 2025), suggesting that Area X may also contain multiple subclasses of GPe-like pallidal neurons. Xiao et al. (2021) reported that a cluster identified as pallidal neurons by snRNA-seq expressed high levels of markers for a type of GPe neuron known as arkypallidal (Hegeman et al., 2016), which projects back to striatum in mammals. In accordance with this idea, there are no pallidal projections from Area X to the subthalamic nucleus (STN), and pallidal neurons within Area X seem to make collateral projections onto spiny neurons (Farries et al., 2005; Gale & Perkel, 2010). Further characterization based on additional molecular markers combined with patterns of connectivity in both juvenile and adult zebra finches is necessary to distinguish whether the non-projecting pallidal cells of CORE and SHELL regions of Area X contain distinct subclasses.

FoxP2 and DARPP-32 may colocalize in a specific subclass of pallidal neurons only in juvenile birds and could represent a transient feature of vocal learning. Dendritic branching and spine density of medium spiny neurons increase in Area X of zebra finches between 45 and 60 dph and are disrupted by experimental decreases in FoxP2 (Jarrell et al., 2025). In accord with this result, the expression of FoxP2 increases within Area X of juvenile zebra finches between 35 and 50 dph (Haesler et al., 2004; Teramitsu et al., 2010), which corresponds to the age range of our juvenile birds (∼45 dph). This age range includes early stages of sensorimotor integration when birds are learning to imitate a memorized tutor song. Increased co-expression of FoxP2 and DARPP-32 may occur within IND pallidal neurons only during this period of vocal learning. Because we did not examine the expression of DARPP-32 in adult birds, this remains an interesting future question. DARPP-32 is a phosphoprotein that is coupled to multiple receptors, including D1/D5 receptors, D2 and adenosine A2a receptors, and NMDA receptors (Blank, 1997; Langley et al., 1997; Yger & Girault, 2011). In striatal tissue, DARPP-32 is necessary for the cAMP-dependent enhancement of NMDA responses (Blank, 1997). Pallidal neurons in Area X are known to receive direct glutamatergic projections from both HVC and LMAN (Farries et al., 2005) as well as from an excitatory local interneuron (Budzillo et al., 2017). Although it is not known whether these excitatory inputs to pallidal neurons include an NMDAR-mediated component, NMDA responses are ubiquitous in projections from DLM to LMAN and from HVC and LMAN to RA (Bottjer, 2005; Bottjer et al., 1998; Mooney & Konishi, 1991; Stark & Perkel, 1999). It is also not known whether pallidal neurons in Area X receive direct dopaminergic inputs, although DA projections to pallidal cells occur in primates and rodents (Eid & Parent, 2016; Gauthier et al., 1999; Lindvall & Bjorklund, 1979). One intriguing possibility is that DARPP-32 may help to regulate interactions between glutamatergic and dopaminergic afferents by regulating the enhancement of NMDA responses (Blank, 1997; Nair et al., 2016).

Overall, our findings indicate that FoxP2 staining, along with size measurements of nuclei and somata, may be useful as a marker to distinguish the indirect pallidal neuron population in Area X that forms close contacts with direct DLM-projecting pallidal neurons (Farries et al., 2005). In terms of functional roles, the direct pathway of the basal ganglia increases thalamic activity, whereas the indirect pathway is thought to oppose the DIR pathway to decrease thalamic activity (Alexander & Crutcher, 1990). When self-generated vocal motor output matches the memorized tutor song, the direct pathway may enhance firing in DLM to generate a positive reinforcement signal for “correct” vocal behavior. In contrast, pallidal neurons that receive inputs from spiny neurons in a “traditional” indirect pathway would suppress firing in DLM. In that case, the indirect pathway may be involved in generating a negative reinforcement signal for “incorrect” behavior when vocalizations differ from memorized tutor sounds. However, because some pallidal neurons in Area X receive direct cortical inputs from HVC and LMAN, those excitatory inputs might increase firing rates in a subset of IND neurons, which would decrease firing rates in DIR neurons and thus increase thalamic activity (Farries et al., 2005). Understanding how these pathways connect and process motor skill information within the LMAN→Area X→DLM→LMAN feedback loop will provide insight into what roles the basal ganglia pathways play in vocal learning in songbirds as well as motor learning in humans.

## Conflicts of Interest

The authors declare no competing financial interests.

## Acknowledgements

This research was supported by NSF grant 1940957. We thank Dr. Jason Junge and Dr. Arkadi Shwartz of the Translational Imaging Center at the USC Michelson Center for Convergent Bioscience for their assistance in training. We thank Dr. Hugh C. Hemmings for the gift of antibody to DARPP-32.

## Notes

### Competing Interest Statement

The authors have declared no competing interest.

### Summary of Updates

Exposition of text has been improved and more appropriate statistical tests have been used.

## REFERENCES

Achiro, J. M., & Bottjer, S. W. (2013). Neural representation of a target auditory memory in a cortico-basal ganglia pathway. J Neurosci, 33(36), 14475–14488. 10.1523/JNEUROSCI.0710-13.2013

Achiro, J. M., Shen, J., & Bottjer, S. W. (2017). Neural activity in cortico-basal ganglia circuits of juvenile songbirds encodes performance during goal-directed learning. Elife, 6. 10.7554/eLife.26973

Al Ghamdi, K. S., Polgar, E., & Todd, A. J. (2009). Soma size distinguishes projection neurons from neurokinin 1 receptor-expressing interneurons in lamina I of the rat lumbar spinal dorsal horn. Neuroscience, 164(4), 1794–1804. 10.1016/j.neuroscience.2009.09.071

Alexander, G. E., & Crutcher, M. D. (1990). Functional architecture of basal ganglia circuits: neural substrates of parallel processing. Trends Neurosci, 13(7), 266–271. http://www.ncbi.nlm.nih.gov/entrez/query.fcgi?cmd=Retrieve&db=PubMed&dopt=Citation&list_uids=1695401

Alvarez-Buylla, A., & Nottebohm, F. (1988). Migration of young neurons in adult avian brain. Nature, 335(6188), 353–354. http://www.ncbi.nlm.nih.gov/entrez/query.fcgi?cmd=Retrieve&db=PubMed&dopt=Citation&list_uids=3419503

Anderson, K. D., & Reiner, A. (1991). Immunohistochemical localization of DARPP-32 in striatal projection neurons and striatal interneurons: implications for the localization of D1-like dopamine receptors on different types of striatal neurons. Brain Resaerch, 568(1-2), 235–243. 10.1016/0006-8993(91)91403-n

Arber, S., & Costa, R. M. (2022). Networking brainstem and basal ganglia circuits for movement. Nat Rev Neurosci, 23(6), 342–360. 10.1038/s41583-022-00581-w

Aronov, D., Andalman, A. S., & Fee, M. S. (2008). A specialized forebrain circuit for vocal babbling in the juvenile songbird. Science, 320(5876), 630–634. 320/5876/630 [pii] 10.1126/science.1155140

Benjamini, Y., & Hochberg, Y. (1995). Controlling the False Discovery Rate - a Practical and Powerful Approach to Multiple Testing. Journal of the Royal Statistical Society Series B-Methodological, 57(1), 289–300. <GO to ISI>://WOS:A1995QE45300017

Blank, R. H. (1997). Assisted reproduction and reproductive rights: the case of in vitro fertilization. Politics Life Sci, 16(2), 279–288. 10.1017/s0730938400024849

Böhner, J. (1983). Song Learning in the Zebra Finch (Taeniopygia-Guttata) - Selectivity in the Choice of a Tutor and Accuracy of Song Copies. Animal Behaviour, 31(Feb), 231–237. 10.1016/S0003-3472(83)80193-6

Böhner, J. (1990). Early acquisition of song in the zebra finch, *Taeniopygia guttata*. Animal Behaviour, 39, 369–374. <GO to ISI>://A1990CM68100020

Bostan, A. C., Dum, R. P., & Strick, P. L. (2018). Functional Anatomy of Basal Ganglia Circuits with the Cerebral Cortex and the Cerebellum. Prog Neurol Surg, 33, 50–61. 10.1159/000480748

Bottjer, S. W. (2005). Silent synapses in a thalamo-cortical circuit necessary for song learning in zebra finches. Journal of Neurophysiology, 94(6), 3698–3707. http://www.ncbi.nlm.nih.gov/entrez/query.fcgi?cmd=Retrieve&db=PubMed&dopt=Citation&list_uids=16107531

Bottjer, S. W., & Altenau, B. (2010). Parallel pathways for vocal learning in basal ganglia of songbirds. Nat Neurosci, 13(2), 153–155. 10.1038/nn.2472

Bottjer, S. W., Brady, J. D., & Cribbs, B. (2000). Connections of a motor cortical region in zebra finches: relation to pathways for vocal learning. J Comp Neurol, 420(2), 244–260. http://www.ncbi.nlm.nih.gov/entrez/query.fcgi?cmd=Retrieve&db=PubMed&dopt=Citation&list_uids=10753310

Bottjer, S. W., Brady, J. D., & Walsh, J. P. (1998). Intrinsic and synaptic properties of neurons in the vocal-control nucleus IMAN from in vitro slice preparations of juvenile and adult zebra finches. J Neurobiol, 37(4), 642–658. http://www.ncbi.nlm.nih.gov/entrez/query.fcgi?cmd=Retrieve&db=PubMed&dopt=Citation&list_uids=9858265

Bottjer, S. W., Halsema, K. A., Brown, S. A., & Miesner, E. A. (1989). Axonal connections of a forebrain nucleus involved with vocal learning in zebra finches. J Comp Neurol, 279(2), 312–326. http://www.ncbi.nlm.nih.gov/entrez/query.fcgi?cmd=Retrieve&db=PubMed&dopt=Citation&list_uids=2464011

Bottjer, S. W., Miesner, E. A., & Arnold, A. P. (1984). Forebrain lesions disrupt development but not maintenance of song in passerine birds. Science, 224(4651), 901–903. http://www.ncbi.nlm.nih.gov/entrez/query.fcgi?cmd=Retrieve&db=PubMed&dopt=Citation&list_uids=6719123

Budzillo, A., Duffy, A., Miller, K. E., Fairhall, A. L., & Perkel, D. J. (2017). Dopaminergic modulation of basal ganglia output through coupled excitation-inhibition. Proceedings of the National Academy of Sciences of the United States of America, 114(22), 5713–5718. 10.1073/pnas.1611146114

Campbell, P., Reep, R. L., Stoll, M. L., Ophir, A. G., & Phelps, S. M. (2009). Conservation and diversity of Foxp2 expression in muroid rodents: functional implications. J Comp Neurol, 512(1), 84–100. 10.1002/cne.21881

Carrillo, G. D., & Doupe, A. J. (2004). Is the songbird Area X striatal, pallidal, or both? An anatomical study. J Comp Neurol, 473(3), 415–437. http://www.ncbi.nlm.nih.gov/entrez/query.fcgi?cmd=Retrieve&db=PubMed&dopt=Citation&list_uids=15116398

Catchpole, C., & Slater, P. J. B. (1995). Bird song: biological themes and variations. New York, NY Cambridge University Press. Publisher description http://www.loc.gov/catdir/description/cam026/94026746.html

Clayton, N. S. (1987). Song Learning in Cross-Fostered Zebra Finches - a Reexamination of the Sensitive Phase. Behaviour, 102, 67–81.

Colquitt, B. M., Merullo, D. P., Konopka, G., Roberts, T. F., & Brainard, M. S. (2021). Cellular transcriptomics reveals evolutionary identities of songbird vocal circuits. Science, 371(6530). 10.1126/science.abd9704

Courtney, C. D., Pamukcu, A., & Chan, C. S. (2023). Cell and circuit complexity of the external globus pallidus. Nat Neurosci, 26(7), 1147–1159. 10.1038/s41593-023-01368-7

Day, N. F., Hobbs, T. G., Heston, J. B., & White, S. A. (2019). Beyond Critical Period Learning: Striatal FoxP2 Affects the Active Maintenance of Learned Vocalizations in Adulthood. eNeuro, 6(2). 10.1523/ENEURO.0071-19.2019

den Hoed, J., & Fisher, S. E. (2020). Genetic pathways involved in human speech disorders. Curr Opin Genet Dev, 65, 103–111. 10.1016/j.gde.2020.05.012

Durstewitz, D., Kroner, S., Hemmings, H. C., Jr., & Gunturkun, O. (1998). The dopaminergic innervation of the pigeon telencephalon: distribution of DARPP-32 and co-occurrence with glutamate decarboxylase and tyrosine hydroxylase. Neuroscience, 83(3), 763–779. 10.1016/s0306-4522(97)00450-8

Eales, L. A. (1985). Song learning in zebra finches - some effects of song model availability on what is learnt and when Animal Behaviour, 33(NOV), 1293–1300. <GO to ISI>://WOS:A1985AUK6100026

Eid, L., & Parent, M. (2016). Chemical anatomy of pallidal afferents in primates. Brain Struct Funct, 221(9), 4291–4317. 10.1007/s00429-016-1216-y

Farries, M. A., Ding, L., & Perkel, D. J. (2005). Evidence for “direct” and “indirect” pathways through the song system basal ganglia. J Comp Neurol, 484(1), 93–104. http://www.ncbi.nlm.nih.gov/entrez/query.fcgi?cmd=Retrieve&db=PubMed&dopt=Citation&list_uids=15717304

Foster, E. F., Mehta, R. P., & Bottjer, S. W. (1997). Axonal connections of the medial magnocellular nucleus of the anterior neostriatum in zebra finches. J Comp Neurol, 382(3), 364–381. http://www.ncbi.nlm.nih.gov/entrez/query.fcgi?cmd=Retrieve&db=PubMed&dopt=Citation&list_uids=9183699

Gale, S. D., & Perkel, D. J. (2010). Anatomy of a songbird basal ganglia circuit essential for vocal learning and plasticity. J Chem Neuroanat, 39(2), 124–131. 10.1016/j.jchemneu.2009.07.003

Gale, S. D., Person, A. L., & Perkel, D. J. (2008). A novel basal ganglia pathway forms a loop linking a vocal learning circuit with its dopaminergic input. J Comp Neurol, 508(5), 824–839. 10.1002/cne.21700

Gauthier, J., Parent, M., Levesque, M., & Parent, A. (1999). The axonal arborization of single nigrostriatal neurons in rats. Brain Resaerch, 834(1-2), 228–232. 10.1016/s0006-8993(99)01573-5

Goldberg, J. H., Adler, A., Bergman, H., & Fee, M. S. (2010). Singing-related neural activity distinguishes two putative pallidal cell types in the songbird basal ganglia: comparison to the primate internal and external pallidal segments. J Neurosci, 30(20), 7088–7098. 10.1523/JNEUROSCI.0168-10.2010

Graybiel, A. M. (2008). Habits, rituals, and the evaluative brain. Annual Review of Neuroscience, 31, 359–387. 10.1146/annurev.neuro.29.051605.112851

Haesler, S., Rochefort, C., Georgi, B., Licznerski, P., Osten, P., & Scharff, C. (2007). Incomplete and inaccurate vocal imitation after knockdown of FoxP2 in songbird basal ganglia nucleus Area X. PLoS Biol, 5(12), e321. 10.1371/journal.pbio.0050321

Haesler, S., Wada, K., Nshdejan, A., Morrisey, E. E., Lints, T., Jarvis, E. D., & Scharff, C. (2004). FoxP2 expression in avian vocal learners and non-learners. J Neurosci, 24(13), 3164–3175. 10.1523/JNEUROSCI.4369-03.2004

Hegeman, D. J., Hong, E. S., Hernandez, V. M., & Chan, C. S. (2016). The external globus pallidus: progress and perspectives. Eur J Neurosci, 43(10), 1239–1265. 10.1111/ejn.13196

Hemmings, H. C., Jr., & Greengard, P. (1986). DARPP-32, a dopamine- and adenosine 3’:5’-monophosphate-regulated phosphoprotein: regional, tissue, and phylogenetic distribution. J Neurosci, 6(5), 1469–1481. 10.1523/JNEUROSCI.06-05-01469.1986

Heston, J. B., & White, S. A. (2015). Behavior-linked FoxP2 regulation enables zebra finch vocal learning. J Neurosci, 35(7), 2885–2894. 10.1523/JNEUROSCI.3715-14.2015

Immelmann, K. (Ed.). (1969). Song development in the zebra finch and other estrildid finches (Vol. Bird Vocalizations) [book chapter].

Iyengar, S., Viswanathan, S. S., & Bottjer, S. W. (1999). Development of topography within song control circuitry of zebra finches during the sensitive period for song learning. J Neurosci, 19(14), 6037–6057. http://www.ncbi.nlm.nih.gov/entrez/query.fcgi?cmd=Retrieve&db=PubMed&dopt=Citation&list_uids=10407041

Jarrell, H., Akhtar, A., Horowitz, M., Huang, Z., Shi, Z., Fang, Z., & Li, X. (2025). Change of Spiny Neuron Structure in the Basal Ganglia Song Circuit and Its Regulation by miR-9 during Song Development. J Neurosci, 45(29). 10.1523/JNEUROSCI.2276-23.2025

Johnson, F., & Bottjer, S. W. (1992). Growth and regression of thalamic efferents in the song-control system of male zebra finches. J Comp Neurol, 326(3), 442–450. http://www.ncbi.nlm.nih.gov/entrez/query.fcgi?cmd=Retrieve&db=PubMed&dopt=Citation&list_uids=1469121

Johnson, F., Sablan, M. M., & Bottjer, S. W. (1995). Topographic organization of a forebrain pathway involved with vocal learning in zebra finches. J Comp Neurol, 358(2), 260–278. http://www.ncbi.nlm.nih.gov/entrez/query.fcgi?cmd=Retrieve&db=PubMed&dopt=Citation&list_uids=7560286

Kosubek-Langer, J., & Scharff, C. (2020). Dynamic FoxP2 levels in male zebra finches are linked to morphology of adult-born Area X medium spiny neurons. Sci Rep, 10(1), 4787. 10.1038/s41598-020-61740-6

Kubikova, L., Wada, K., & Jarvis, E. D. (2010). Dopamine receptors in a songbird brain. J Comp Neurol, 518(6), 741–769. 10.1002/cne.22255

Langley, K. C., Bergson, C., Greengard, P., & Ouimet, C. C. (1997). Co-localization of the D1 dopamine receptor in a subset of DARPP-32-containing neurons in rat caudate-putamen. Neuroscience, 78(4), 977–983. 10.1016/s0306-4522(96)00583-0

Lindvall, O., & Bjorklund, A. (1979). Dopaminergic innervation of the globus pallidus by collaterals from the nigrostriatal pathway. Brain Resaerch, 172(1), 169–173. 10.1016/0006-8993(79)90907-7

Luo, M., Ding, L., & Perkel, D. J. (2001). An avian basal ganglia pathway essential for vocal learning forms a closed topographic loop. J Neurosci, 21(17), 6836–6845. http://www.ncbi.nlm.nih.gov/entrez/query.fcgi?cmd=Retrieve&db=PubMed&dopt=Citation&list_uids=11517271

Luo, M., & Perkel, D. J. (1999a). A GABAergic, strongly inhibitory projection to a thalamic nucleus in the zebra finch song system. J Neurosci, 19(15), 6700–6711. http://www.ncbi.nlm.nih.gov/entrez/query.fcgi?cmd=Retrieve&db=PubMed&dopt=Citation&list_uids=10414999

Luo, M., & Perkel, D. J. (1999b). Long-range GABAergic projection in a circuit essential for vocal learning. J Comp Neurol, 403(1), 68–84. http://www.ncbi.nlm.nih.gov/entrez/query.fcgi?cmd=Retrieve&db=PubMed&dopt=Citation&list_uids=10075444

Mann, N. I., & Slater, P. J. B. (1995). Song Tutor Choice by Zebra Finches in Aviaries. Animal Behaviour, 49(3), 811–820. <GO to ISI>://WOS:A1995QM72800023

Mastro, K. J., Zitelli, K. T., Willard, A. M., Leblanc, K. H., Kravitz, A. V., & Gittis, A. H. (2017). Cell-specific pallidal intervention induces long-lasting motor recovery in dopamine-depleted mice. Nat Neurosci, 20(6), 815–823. 10.1038/nn.4559

Mooney, R., & Konishi, M. (1991). Two distinct inputs to an avian song nucleu activate different glutamate receptor subtypes on individual neurons Proceedings of the National Academy of Sciences of the United States of America, 88(10), 4075–4079. <GO to ISI>://WOS:A1991FM04200004

Murugan, M., Harward, S., Scharff, C., & Mooney, R. (2013). Diminished FoxP2 levels affect dopaminergic modulation of corticostriatal signaling important to song variability. Neuron, 80(6), 1464–1476. 10.1016/j.neuron.2013.09.021

Nair, A. G., Bhalla, U. S., & Hellgren Kotaleski, J. (2016). Role of DARPP-32 and ARPP-21 in the Emergence of Temporal Constraints on Striatal Calcium and Dopamine Integration. PLoS Comput Biol, 12(9), e1005080. 10.1371/journal.pcbi.1005080

Nicholson, D. A., Roberts, T. F., & Sober, S. J. (2018). Thalamostriatal and cerebellothalamic pathways in a songbird, the Bengalese finch. J Comp Neurol, 526(9), 1550–1570. 10.1002/cne.24428

Nixdorf-Bergweiler, B. E., Lips, M. B., & Heinemann, U. (1995). Electrophysiological and morphological evidence for a new projection of LMAN-neurones towards area X. Neuroreport, 6(13), 1729–1732. http://www.ncbi.nlm.nih.gov/entrez/query.fcgi?cmd=Retrieve&db=PubMed&dopt=Citation&list_uids=8541469

Nolan, M., Scott, C., Hof, P. R., & Ansorge, O. (2024). Betz cells of the primary motor cortex. J Comp Neurol, 532(1), e25567. 10.1002/cne.25567

Nottebohm, F., Stokes, T. M., & Leonard, C. M. (1976). Central control of song in the canary, Serinus canarius. J Comp Neurol, 165(4), 457–486. http://www.ncbi.nlm.nih.gov/entrez/query.fcgi?cmd=Retrieve&db=PubMed&dopt=Citation&list_uids=1262540

Oorschot, D. E. (2016). Cell types in the different nuclei of the basal ganglia In Handbook of Behavioral Neuroscience. (Vol. 24, pp. 99–117.). Elsevier. 10.1016/B978-0-12-802206-1.00005-2

Ouimet, C. C., Langley-Gullion, K. C., & Greengard, P. (1998). Quantitative immunocytochemistry of DARPP-32-expressing neurons in the rat caudatoputamen. Brain Resaerch, 808(1), 8–12. 10.1016/s0006-8993(98)00724-0

Paterson, A. K., & Bottjer, S. W. (2017). Cortical inter-hemispheric circuits for multimodal vocal learning in songbirds. J Comp Neurol, 525(15), 3312–3340. 10.1002/cne.24280

Person, A. L., Gale, S. D., Farries, M. A., & Perkel, D. J. (2008). Organization of the songbird basal ganglia, including area X. J Comp Neurol, 508(5), 840–866. 10.1002/cne.21699

Pidoux, L., Le Blanc, P., Levenes, C., & Leblois, A. (2018). A subcortical circuit linking the cerebellum to the basal ganglia engaged in vocal learning. Elife, 7. 10.7554/eLife.32167

Pinaud, R., Saldanha, C. J., Wynne, R. D., Lovell, P. V., & Mello, C. V. (2007). The excitatory thalamo-“cortical” projection within the song control system of zebra finches is formed by calbindin-expressing neurons. J Comp Neurol, 504(6), 601–618. http://www.ncbi.nlm.nih.gov/entrez/query.fcgi?cmd=Retrieve&db=PubMed&dopt=Citation&list_uids=17722049

Redgrave, P., Rodriguez, M., Smith, Y., Rodriguez-Oroz, M. C., Lehericy, S., Bergman, H., Agid, Y., DeLong, M. R., & Obeso, J. A. (2010). Goal-directed and habitual control in the basal ganglia: implications for Parkinson’s disease. Nat Rev Neurosci, 11(11), 760–772. 10.1038/nrn2915

Reiner, A., Laverghetta, A. V., Meade, C. A., Cuthbertson, S. L., & Bottjer, S. W. (2004). An immunohistochemical and pathway tracing study of the striatopallidal organization of area X in the male zebra finch. J Comp Neurol, 469(2), 239–261. http://www.ncbi.nlm.nih.gov/entrez/query.fcgi?cmd=Retrieve&db=PubMed&dopt=Citation&list_uids=14694537

Reiner, A., Perera, M., Paullus, R., & Medina, L. (1998). Immunohistochemical localization of DARPP32 in striatal projection neurons and striatal interneurons in pigeons. J Chem Neuroanat, 16(1), 17–33. 10.1016/s0891-0618(98)00056-8

Roper, A., & Zann, R. (2006). The onset of song learning and song tutor selection in fledgling zebra finches. Ethology, 112(5), 458–470. 10.1111/j.1439-0310.2005.01169.x

Sanchez-Valpuesta, M., Suzuki, Y., Shibata, Y., Toji, N., Ji, Y., Afrin, N., Asogwa, C. N., Kojima, I., Mizuguchi, D., Kojima, S., Okanoya, K., Okado, H., Kobayashi, K., & Wada, K. (2019). Corticobasal ganglia projecting neurons are required for juvenile vocal learning but not for adult vocal plasticity in songbirds. Proceedings of the National Academy of Sciences of the United States of America, 116(45), 22833–22843. 10.1073/pnas.1913575116

Scharff, C., & Nottebohm, F. (1991). A comparative study of the behavioral deficits following lesions of various parts of the zebra finch song system: implications for vocal learning. J Neurosci, 11(9), 2896–2913. http://www.ncbi.nlm.nih.gov/entrez/query.fcgi?cmd=Retrieve&db=PubMed&dopt=Citation&list_uids=1880555

Shi, Z., Piccus, Z., Zhang, X., Yang, H., Jarrell, H., Ding, Y., Teng, Z., Tchernichovski, O., & Li, X. (2018). miR-9 regulates basal ganglia-dependent developmental vocal learning and adult vocal performance in songbirds. Elife, 7. 10.7554/eLife.29087

Singh, U. A., & Iyengar, S. (2019). The expression of DARPP-32 in adult male zebra finches (Taenopygia guttata). Brain Struct Funct, 224(8), 2939–2972. 10.1007/s00429-019-01947-0

Soares-Cunha, C., Coimbra, B., Sousa, N., & Rodrigues, A. J. (2016). Reappraising striatal D1- and D2-neurons in reward and aversion. Neurosci Biobehav Rev, 68, 370–386. 10.1016/j.neubiorev.2016.05.021

Sohrabji, F., Nordeen, E. J., & Nordeen, K. W. (1990). Selective impairment of song learning following lesions of a forebrain nucleus in the juvenile zebra finch. Behav Neural Biol, 53(1), 51–63. http://www.ncbi.nlm.nih.gov/entrez/query.fcgi?cmd=Retrieve&db=PubMed&dopt=Citation&list_uids=2302141

Spix, T. A., Nanivadekar, S., Toong, N., Kaplow, I. M., Isett, B. R., Goksen, Y., Pfenning, A. R., & Gittis, A. H. (2021). Population-specific neuromodulation prolongs therapeutic benefits of deep brain stimulation. Science, 374(6564), 201–206. 10.1126/science.abi7852

Stark, L. L., & Perkel, D. J. (1999). Two-stage, input-specific synaptic maturation in a nucleus essential for vocal production in the zebra finch. J Neurosci, 19(20), 9107–9116. http://www.ncbi.nlm.nih.gov/entrez/query.fcgi?cmd=Retrieve&db=PubMed&dopt=Citation&list_uids=10516328

Teramitsu, I., Poopatanapong, A., Torrisi, S., & White, S. A. (2010). Striatal FoxP2 is actively regulated during songbird sensorimotor learning. PLoS ONE, 5(1), e8548. 10.1371/journal.pone.0008548

Thompson, C. K., Schwabe, F., Schoof, A., Mendoza, E., Gampe, J., Rochefort, C., & Scharff, C. (2013). Young and intense: FoxP2 immunoreactivity in Area X varies with age, song stereotypy, and singing in male zebra finches. Front Neural Circuits, 7, 24. 10.3389/fncir.2013.00024

Tomasi, S., Caminiti, R., & Innocenti, G. M. (2012). Areal differences in diameter and length of corticofugal projections. Cereb Cortex, 22(6), 1463–1472. 10.1093/cercor/bhs011

Trusel, M., Zhao, Z., Alam, D. H., Marks, E. S., Ikeda, M. Z., & Roberts, T. F. (2025). Synaptic connectivity of sensorimotor circuits for vocal imitation in the songbird. Elife, 14. 10.7554/eLife.104609

Vates, G. E., & Nottebohm, F. (1995). Feedback circuitry within a song-learning pathway. Proceedings of the National Academy of Sciences of the United States of America, 92(11), 5139–5143. http://www.ncbi.nlm.nih.gov/entrez/query.fcgi?cmd=Retrieve&db=PubMed&dopt=Citation&list_uids=7761463

Vounatsos, M. V., & Gittis, A. H. (2025). GPe Projections to the Retrorubral Field Give Rise to Diverging GABAergic and DAergic Circuits. J Neurosci, 45(2). 10.1523/JNEUROSCI.1446-24.2024

Xiao, L., Merullo, D. P., Koch, T. M. I., Cao, M., Co, M., Kulkarni, A., Konopka, G., & Roberts, T. F. (2021). Expression of FoxP2 in the basal ganglia regulates vocal motor sequences in the adult songbird. Nat Commun, 12(1), 2617. 10.1038/s41467-021-22918-2

Yger, M., & Girault, J. A. (2011). DARPP-32, Jack of All Trades… Master of Which? Front Behav Neurosci, 5, 56. 10.3389/fnbeh.2011.00056

Yin, H. H., & Knowlton, B. J. (2006). The role of the basal ganglia in habit formation. Nat Rev Neurosci, 7(6), 464–476. 10.1038/nrn1919

Zaremba, B., Fallahshahroudi, A., Schneider, C., Schmidt, J., Sarropoulos, I., Leushkin, E., Berki, B., Van Poucke, E., Jensen, P., Senovilla-Ganzo, R., Hervas-Sotomayor, F., Trost, N., Lamanna, F., Sepp, M., Garcia-Moreno, F., & Kaessmann, H. (2025). Developmental origins and evolution of pallial cell types and structures in birds. Science, 387(6735), eadp5182. 10.1126/science.adp5182

